# Geminivirus Replication Protein impairs SUMO conjugation of PCNA at two acceptor sites

**DOI:** 10.1101/305789

**Authors:** Manuel Arroyo-Mateos, Blanca Sabarit, Francesca Maio, Miguel A. Sánchez-Durán, Tabata Rosas-Díaz, Marcel Prins, Javier Ruiz-Albert, Ana P Luna, Harrold A. van den Burg, Eduardo R. Bejarano

**Affiliations:** Instituto de Hortofruticultura Subtropical y Mediterránea “La Mayora”, Universidad de Málaga-Consejo Superior de Investigaciones Científicas (IHSM-UMA-CSIC), Dept. Biología Celular, Genética y Fisiología, Universidad de Málaga, Campus Teatinos, 29071 Málaga, Spain.; Molecular Plant Pathology, Swammerdam Institute for Life Sciences, Faculty of Science University of Amsterdam, PO box 94215,1090 GE Amsterdam, The Netherlands.; Keygene NV, Wageningen, The Netherlands.

**Keywords:** Geminivirus, Rep, PCNA, begomovirus, SUMO, sumoylation, homologous recombination

## Abstract

Geminiviruses are DNA viruses that replicate in nuclei of infected plant cells using the plant DNA replication machinery, including PCNA (Proliferating cellular nuclear antigen), a cofactor that orchestrates genome duplication and maintenance by recruiting crucial players to replication forks. These viruses encode a multifunctional protein, Rep, which is essential for viral replication, induces the accumulation of the host replication machinery and interacts with several host proteins, including PCNA and the SUMO E2 conjugation enzyme (SCE1). Post-translational modification of PCNA by ubiquitin or SUMO plays an essential role in the switching of PCNA between interacting partners during DNA metabolism processes (e.g. replication, recombination, repair, etc.). In yeast, PCNA sumoylation has been associated to DNA repair involving homologous recombination (HR). Previously, we reported that ectopic Rep expression results in very specific changes in the sumoylation pattern of plant cells. In this work, we show, using a reconstituted sumoylation system in *Escherichia coli*, that tomato PCNA is sumoylated at two residues, K254 and K164, and that co-expression of the geminivirus protein Rep suppresses sumoylation at these lysines. Finally, we confirm that PCNA is sumoylated *in planta* and that Rep also interferes with PCNA sumoylation in plant cells.

**Importance:** SUMO adducts have a key role in regulating the activity of animal and yeast PCNA on DNA repair and replication. Our work demonstrates for the first time that sumoylation of plant PCNA occurs in plant cells and that a plant virus interferes with this modification. This work marks the importance of sumoylation in allowing viral infection and replication in plants. Moreover, it constitutes a prime example of viral proteins interfering with post-translational modifications of selected host factors to create a proper environment for infection.

## INTRODUCTION

Geminiviruses constitute a large family of plant viruses with circular singlestranded (ss) DNA genomes packaged within geminate particles (1), which replicate in the nuclei of infected cells through double-stranded (ds) DNA intermediates (2, 3). The largest geminivirus genus corresponds to begomoviruses, which can have bipartite genomes (A and B components) like *Tomato golden mosaic virus* (TGMV) or monopartite genomes like *Tomato yellow leaf curl virus* (TYLCV). Begomoviruses encode two proteins involved in viral replication: Rep (also called AL1, AC1, and C1), a multifunctional essential protein, and C3 (also called AL3, AC3, C3, and REn), which interacts with Rep and greatly enhances begomovirus DNA accumulation in host cells (4). Rep has different functions: it mediates recognition of its cognate origin-of-replication in a geminivirus species-specific manner (5), it is required for initiation and termination of viral DNA synthesis (6–8), and it acts as a DNA helicase (9, 10).

Growing evidence strongly supports the notion that geminivirus proteins have a significant impact on a variety of host processes including cell differentiation, cell cycle control, DNA replication, plasmodesmata function and RNA silencing (3). By these means, geminiviruses reshape their environment by co-opting cellular processes necessary for viral replication, systemic spread, and impairment of plant defences. There are numerous mechanisms by which geminiviruses mediate their effects on the host cell, including targeting of post-translational modification systems. Such systems play critical roles in many cellular processes because they cause rapid changes in (i) the function of preexisting proteins, (ii) composition of multi-protein complexes and (iii) their subcellular localization. Their versatility in regulating protein function and cellular behaviour makes them a particularly attractive target for viruses. One example of a key cellular regulatory system targeted by viruses is sumoylation (11, 12), a post-translational process mainly involved in nuclear functions that modifies protein function, activity or localization of its targets through covalent attachment of a 10-kD Ubiquitin-like polypeptide called SUMO (Small ubiquitin-like modifier) (13–15).

Briefly, post-translational modification by SUMO involves a cascade of ATP-dependent reactions that are mechanistically similar to ubiquitination, involving sequential activation and conjugation of SUMO. SUMO activation is driven by an E1 enzyme (SUMO-activating enzyme SAE1/SAE2 heterodimer), while SUMO conjugation is mediated by a single E2 enzyme (SUMO-conjugating enzyme SCE1, also known as Ubc9 in yeast and mammals). The final transfer of SUMO from SCE1 to specific lysine residues in target proteins can occur directly or can be enhanced by SUMO ligases (14, 16). Target proteins can undergo mono-sumoylation of one lysine, poly-sumoylation (SUMO chain formation) or multi-sumoylation (modification of several lysines in one substrate) (17–19). SUMO can be specifically detached from modified lysines by SUMO proteases (ULPs), making it a reversible and dynamic process (18, 20). The consequence of sumoylation on targets is very diverse, ranging from changes in localization to altered activity and, in some cases, stabilization of the modified protein. All of these effects are frequently the result of changes in the molecular interactions of the sumoylated proteins. Sumoylation can either mask a binding site in its target thus inhibiting its interactions with other proteins, increase the number of binding sites on its target hence facilitating the binding of molecules, such as proteins or DNA, or produce a conformational change that modulates its activity.

In plants, the characterization of the sumoylation enzymes has largely been restricted to *Arabidopsis thaliana*, although information based on sequence analysis of other plant genomes is available (21, 22). The *Arabidopsis* genome encodes eight full-length SUMO genes (*AtSUMOs*), a single gene encoding the SUMO-conjugating enzyme SCE1 (*AtSCE1a*) and a large number of ULPs. Only two SUMO E3 ligases (SIZ1, HPY2/MMS21) have been identified and characterized in *Arabidopsis* (23–27). In plants, sumoylation is important for embryonic development, organ growth, flowering transition, and hormone regulation (28). In addition, SUMO also plays a key role in stress-associated responses to stimuli such as extreme temperatures, drought, salinity and nutrient assimilation (29, 30). During these abiotic stresses, the profile of SUMO-modified proteins changes dramatically, greatly increasing the global SUMO-conjugates levels and decreasing the pool of free SUMO (31, 32). After exertion of stress, SUMO-conjugates slowly diminish by the action of ULPs, that are fundamental players in fine-tuning SUMO conjugation/deconjugation (20, 33). Several observations, including pathogen manipulation of SUMO conjugation by bacterial elicitors (34–36), modification of SUMO levels altering pathogen infection in plants (37, 38) and sumoylation influencing innate immunity (39–41), indicate that SUMO also plays an important role in plant defence responses.

Numerous studies in recent years have shown that sumoylation also plays a role in viral infection. In animals, proteins from DNA and RNA virus families were shown to be sumoylated, and this modification seems to be important for their function. Conversely, proteins encoded by DNA viruses can modify host sumoylation, globally or to certain specific substrates, altering the host environment to facilitate viral replication or to overcome host defences, either by preventing *de novo* sumoylation or by enhancing desumoylation (11, 12, 42, 43).

In sharp contrast with these animal pathosystems, only two examples of an interaction between viral proteins and the sumoylation machinery have been described for plants so far. The only RNA-dependent RNA polymerase of the potyvirus *Turnip mosaic virus* (TuMV), Nlb, is sumoylated and interacts with SUMO3 from *Arabidopsis* (44). Knockout or overexpression of SUMO3 suppresses TuMV replication and attenuates the viral symptoms (45). The other example is the interaction between the begomovirus protein Rep and SCE1 (37). This interaction is essential for viral infection, since Rep mutants impaired in SCE1-binding and plants with altered SUMO levels, showed reduced viral replication (37, 46). Transient expression of Rep in *Nicotiana benthamiana* showed that the interaction between Rep and SCE1 does not alter the global sumoylation pattern *in planta*, but rather may specifically influence SUMO conjugation of a selected subset of host proteins (46).

In this study, we identify PCNA (Proliferating cell nuclear antigen) as such a plant protein whose sumoylation is altered in the presence of the begomovirus protein Rep. Using a reconstituted sumoylation assay in *Escherichia coli*, we demonstrate that tomato PCNA is readily sumoylated at two different lysines (K164 and K254). However, in the presence of Rep, SUMO attachment is compromised at both these acceptor sites. This interference is specific for PCNA, since Rep does not alter sumoylation of a control protein. It does also not depend on the physical interaction between Rep and SCE1. Finally, we are able to detect for the first time sumoylation of PCNA *in planta* and show that the reduction of PCNA sumoylation exerted by Rep also occurs in plant tissues.

## RESULTS

### Rep modulates the sumoylation of PCNA

Ectopic Rep expression alters the sumoylation status of specific host proteins (46). Although full determination of the number and identity of these plant targets will require a comprehensive proteomic analysis, proteins that both interact with Rep and are known to be sumoylated are primary candidates. One of such proteins is PCNA, since its sumoylation has been described for PCNA homologues of yeast, *Xenopus laevis*, mammalians and *Arabidopsis* (47–53) and tomato PCNA binds Rep from several begomoviruses (54, 55).

Sumoylation of *Arabidopsis* PCNA has been described for both homologues encoded in its genome (*AtPCNA1* and *AtPCNA2*) using a sumoylation system reconstituted *in bacteria* (52, 53). In order to assess if tomato PCNA (PCNA) is also sumoylated and if Rep then interferes with its sumoylation, we followed a similar strategy and performed sumoylation assays in *E. coli* as described by (56). In this assay, all components of the sumoylation pathway (SAE1/2, SCE1, and SUMO1) are expressed in *E. coli* together with a potential substrate protein using an inducible system based on T7 promoters, where SAE1, SUMO1 and the target protein are expressed as His-tagged proteins to facilitate protein detection. Three compatible plasmids were co-maintained in *E. coli* in this assay to simultaneously express: (i) mammalian His-SAE1, SAE2 and SCE1 (Ubc9) from a polycistronic RNA, (ii) mammalian His-SUMO1, (iii) His-PCNA from tomato, or alternatively His-PCNA and Rep as a polycistronic mRNA. Protein expression was induced in cells co-transformed with the appropriate plasmids and total protein extracts from these cells were analysed with western blots probed with anti-PCNA, anti-SUMO1/2, anti-His or anti-Rep antibodies (Fig. 1). Cells expressing only PCNA displayed a band of the expected size (32 kD) when the blot was incubated with anti-PCNA (Fig. 1, lane 1). An additional band (PCNA-SUMO) of approximately 55 kD was detected when PCNA was expressed together with the complete sumoylation machinery (E1/E2 and SUMO) (Fig.1, lane 5), but not when it was co-expressed only with the sumoylation machinery without SUMO (Fig. 1 lane 4). Since the apparent mass of this band matches the expected mass for a PCNA-SUMO dimer and a similar band was detected when the blot was incubated with anti-SUMO or anti-His antibodies, we conclude that this 55 kD band corresponds to mono-sumoylated form of PCNA. This result indicates that tomato PCNA, like its yeast, mammalian and *Arabidopsis* homologues, can be sumoylated. The extra protein bands observed on the membrane by the anti-SUMO antibody are similar to those reported previously (56) and likely correspond to free SUMO (SUMO) or SUMO chains (dimers, trimers, etc.). Similar bands were detected with the anti-His antibody.

**Figure 1.**
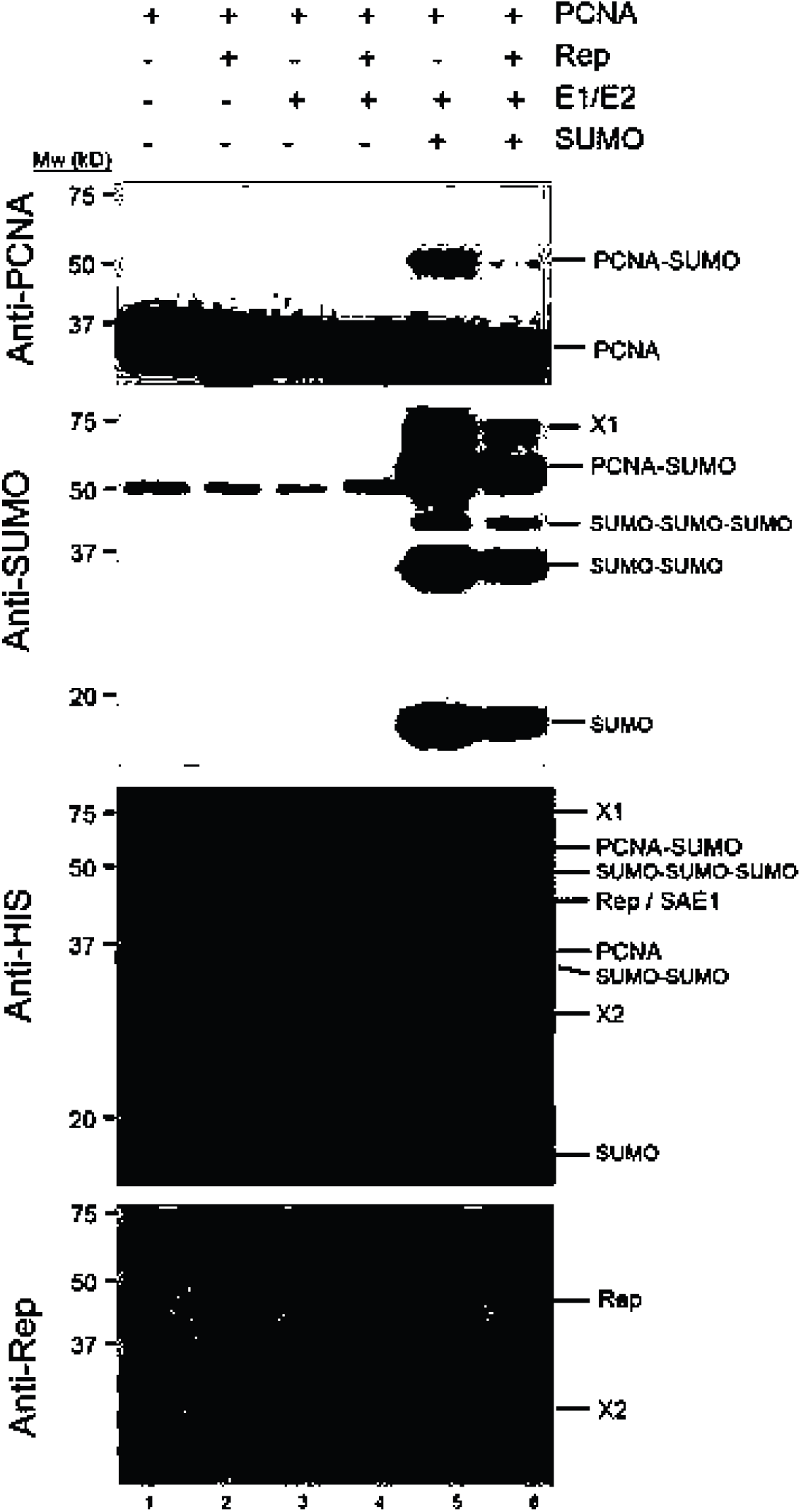
Sumoylation of tomato PCNA in a reconstituted SUMO conjugation system *in bacteria* is modulated by Rep. Tomato PCNA, Rep from *Tomato golden mosaic virus* (TGMV), E1/E2 (mammalian SAE1, SAE2 and Ubc9), SUMO (*HsSUMO1*) were co-expressed in *E. coli* NCM631 cells and extracted. Protein extracts were blotted with the antibodies indicated (left side). Expression (+/-) of the individual components is indicated at the top per lane. Relevant bands are labelled on the right side. The band labelled ‘X1’ could correspond to a SUMO tetramer or more likely to a complex of PCNA-2xSUMO. Band ‘X2’ corresponds to a truncated form of Rep protein. Molecular weight markers (Mw) are indicated.

When PCNA and Rep were co-expressed from the same plasmid as a polycistronic mRNA, together with the complete sumoylation machinery expressed from two accompanying plasmids, the intensity of the band that corresponds to sumoylated PCNA (PCNA-SUMO) was drastically reduced (Fig. 1, lane 6). This reduction in intensity was also observed for a higher molecular weight band (X1) detected by the anti-SUMO antibody. The size of this band is consistent with PCNA with two SUMO peptides attached. However, this extra band was not detected with anti-PCNA, arguing that it reflects a different protein (complex). Together, these results demonstrate that sumoylation of tomato PCNA is strongly reduced in the presence of Rep. When using an anti-Rep antibody, a band corresponding to the expected molecular weight (44 kD) of the viral protein Rep was detected (Fig. 1, anti-Rep). We did not detect any bands that could match with sumoylated forms of Rep, confirming that SUMO is not covalently attached to Rep. The additional smaller band (X2) detected with the anti-Rep antibody likely corresponds to a truncated form of the viral protein. To establish if impairment of PCNA sumoylation is strictly due to co-expression of Rep, we performed a sumoylation assay in which we replaced Rep by the C2 protein from the begomovirus *Tomato yellow leaf curl Sardinia virus*. The results showed that only Rep and not C2 was capable of reducing the amount of sumoylated PCNA, thus ruling out an unspecific interference with sumoylation due to the simultaneous expression of PCNA and any given protein from a polycistronic RNA (Fig. 2A, lanes 7 to 9). Expression of C2 was confirmed using an anti-His antibody.

**Figure 2.**
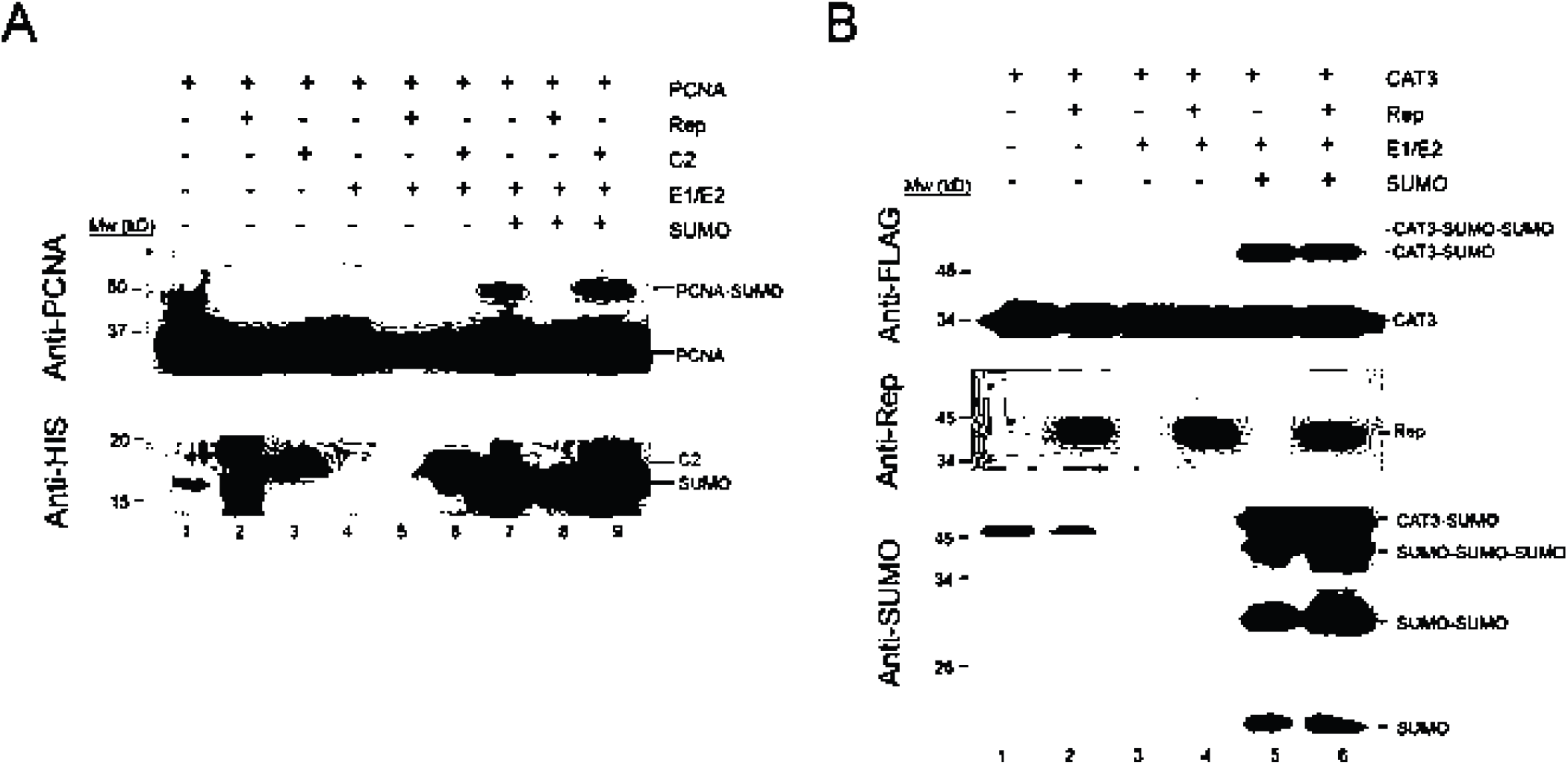
Rep interferes only with sumoylation of PCNA but not with other proteins. (A) Tomato PCNA, Rep from *Tomato golden mosaic virus* (TGMV), C2 from *Tomato yellow leaf curl Sardinia virus* (TYLCSV), E1/E2 (mammalian SAE1, SAE2 and Ubc9), SUMO (*HsSUMO1*) were co-expressed in *E. coli* NCM631 cells and extracted. Protein extracts were blotted with the antibodies indicated antibodies indicated (left side). Expression (+/-) of the individual components is indicated. Relevant bands are labelled on the right side. (B) Similar to (A), except that Catalse3 (CAT3) is co-expressed.

To determine if the Rep-mediated suppression of PCNA sumoylation is due to the inhibition of SCE1 activity by the viral protein, we carried out a sumoylation assay replacing PCNA with *Arabidopsis* Catalase 3 (CAT3), a known sumoylation substrate (57). A C-terminal CAT3 fragment fused to a Flag epitope was expressed in *E. coli* cells in the presence or absence of Rep. Again, western blots were probed with anti-Flag, anti-Rep and anti-SUMO antibodies (Figure 2B). In all proteins extracts, a band of approximately 35 kD corresponding to the CAT3 protein was detected with the anti-Flag antibody (Fig. 2B, anti-FLAG). When CAT3 was co-expressed with the complete sumoylation machinery (E1/E2 and SUMO), an additional band of approximately 50 kD, consistent with mono-sumoylated CAT3, was detected both with the anti-Flag (Fig. 2B, anti-FLAG, lane 5) and the anti-SUMO antibody (Fig. 2B, anti-SUMO, lane 5). When CAT3 and Rep were expressed simultaneously from one plasmid as a polycistronic mRNA, together with the sumoylation machinery expressed from the two accompanying plasmids, the intensity of the band identified as sumoylated CAT3 remained unaltered (Fig. 2B, lane 6). This indicates that Rep expression does not affect the conjugating activity of SCE1 in the *E. coli* assay in a generic way. Rep expression was confirmed by probing the western blot with an anti-Rep antibody.

### Tomato PCNA is sumoylated at the residues K164 and K254

Sumoylation of yeast PCNA occurs preferentially at K164, a residue conserved in all PCNA proteins, and to a lesser extent at K127 (47). Sumoylation of yeast PCNA K164 requires the SUMO E3 *Sc*Siz1, whereas K127 sumoylation proceeds without an E3 ligase *in vitro* and is mediated by *Sc*Siz2 *in vivo* (47, 58). Sumoylation at K164 has been observed in other species, such as chicken cells, *X. laevis* egg extracts and mammalian cells (48–51, 59). In the in *E. coli* reconstituted sumoylation, *Arabidopsis* PCNA was shown to be sumoylated primarily at K254, although additional sumoylation was reported to occur at other lysine residues (K13, K14, K20, K217, and K240), but not at K164 (52).

In order to establish whether tomato PCNA also contains multiple SUMO acceptor sites and to examine if Rep interferes with sumoylation at each site, we performed sumoylation assays expressing tomato PCNA mutants where lysines were replaced by alanines. To select PCNA lysine residues for mutagenesis, we analysed a PCNA multisequence alignment and candidate lysines were picked according to the following criteria: (*i*) the residue is conserved in PCNA homologues of different organisms, (*ii*) the residue was previously described as SUMO acceptor site in other PCNA homologues, (*iii*) the residue is located in a predicted SUMO acceptor site using GPS SUMO-gp and SUMOsp2.0 (60, 61) and/or (*iv*) the residue is located at the surface of the PCNA three-dimensional structure. These analyses suggested the residues K91, K164, K168, K190 and K254 to have increased probability to be sumoylated, with K164 and K254 being the prime candidates (Fig. 3A). As a first step, single and double mutants were generated for the residues K164 and K254 and analysed in our sumoylation assay in *E. coli*. Co-expression of wild-type PCNA with the complete sumoylation machinery (E1/E2 and SUMO) produced a double band (PCNA-SUMO) of the expected molecular mass of sumoylated PCNA (Fig. 3B, lane 2): we did not observe this double band before, due to a lower resolution in protein separation in our previous western blots (Fig. 1). This double band could represent PCNA monomers mono-sumoylated at two different positions. This phenomenon was previously described for yeast and human PCNA, which when sumoylated at K127 or K254 respectively migrated on SDS-PAGE gel at a different apparent molecular mass than yeast PCNA SUMO-modified at K164 (47, 50, 62). When PCNA K164A was co-expressed instead of wild-type PCNA, a single band of 55 kD (corresponding with the lower of the two bands obtained with the wild-type PCNA) was observed (Fig. 3B, lane 4). On the contrary, co-expression of the single PCNA mutant K254A resulted in a decrease in intensity of the lower band, while the intensity of the upper band remained unchanged at the levels seen for wild-type PCNA (Fig. 3B, lane 6). When the PCNA double mutant K164A/K254A was expressed, the upper band disappeared entirely while a decrease in the intensity was detected for the lower band, similar to situation shown for the single mutant K254A (Fig. 3B, lane 5). These results indicated that the double band detected with wild-type PCNA corresponds to two distinct PCNA-SUMO adducts in which SUMO is attached to two different lysines, with the upper band, being the result of sumoylation at K164 and the lower, at least partially, the product of sumoylation at K254. To identify the alternative lysine residue(s) responsible for the residual sumoylation of the lower band in PCNA K254A, we performed sumoylation assays introducing into the PCNA double mutant K164A/K254A additional mutations in each of the other selected lysine residues (K91, K168 or K190). However, none of the additional mutations was able to remove the weak band present in the double mutant K164A/K254A, suggesting that lysine residues other than K91, K168 or K190 are sumoylated *in bacteria* when K164 and K254 are mutated (results with the additional K91A, K168A and K190A mutations are shown in Fig. S1). When co-expressing Rep no lower band is detected and a large reduction in the upper band is observed (Fig. 3B, Lane 3) indicating that expression of Rep impairs PCNA sumoylation in all lysines residues.

**Figure 3.**
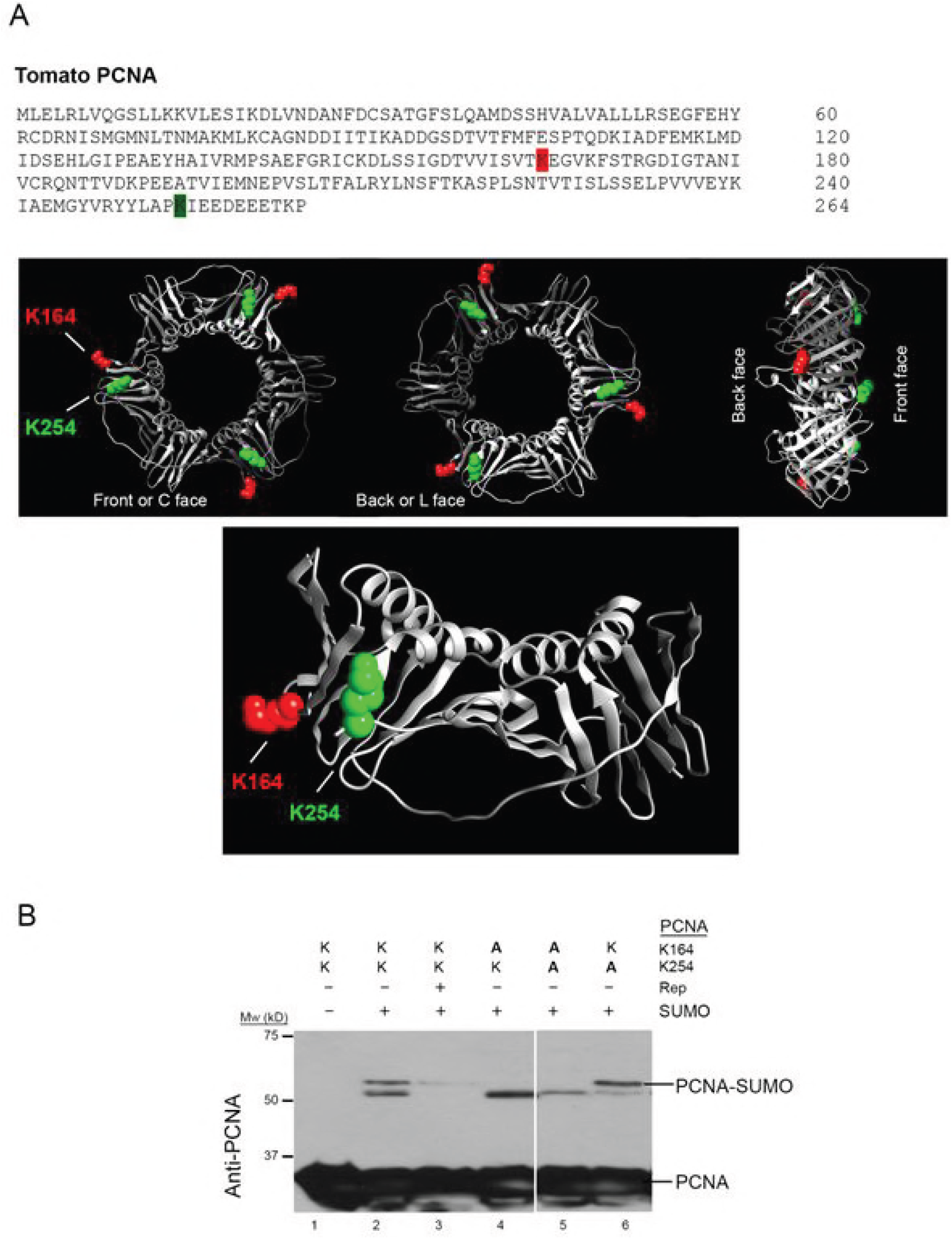
Identification of tomato PCNA SUMO acceptor sites. (A) Primary sequence of tomato PCNA with residues K164 and K254 highlighted (red and green, respectively) (upper panel) and the predicted 3D model of the structure of a tomato PCNA trimer (central panel) or a monomer (lower panel) (B) Sumoylation assay of tomato PCNA single mutants K164A, K254A and the double mutant K164A/K254A *in bacteria* while co-expressing Rep from TGMV. Assay is similar to Figure 1. Expression (+/-) of the individual components is indicated. E1/E2 enzymes were expressed in all samples. Coomassie staining to confirm equal protein loading are showed.

### Rep-mediated suppression of PCNA sumoylation is independent of the Rep-SCE1 interaction

To gain insight into the mechanism of Rep-mediated suppression of PCNA sumoylation, we analysed whether the physical interaction of Rep with SCE1 or PCNA had a role. Previous work had mapped the SCE1-binding domain of Rep between the residues 56-114, with the region 56-85 likely forming the core of the interface with *N. benthamina* SCE1, while the region 86-114 may stabilize or enhance this interaction. Replacement of the lysine residues K68 and K102 in the binding region of Rep impairs its interaction with SCE1 and dramatically reduces viral replication, indicating that the Rep-SCE1 interaction is required for viral DNA replication (37, 46). To determine whether the Rep-SCE1 interaction (37, 46, 54) is required for compromised PCNA sumoylation, we performed a sumoylation assay in *E. coli* co-expressing the Rep K68A/K102A double mutant, which does not interact with *N. benthamiana* SCE1 (46). As expected, the double band corresponding with sumoylated PCNA vanished when wild type Rep was coexpressed (Fig. 4, lane 2). A similar reduction in the intensity of these two bands was seen when Rep K68A/K102A was co-expressed (Fig. 4, lane 3). This indicates that, in the conditions of our sumoylation assay, the direct interaction SCE1-Rep is not required for the Rep-mediated suppression of PCNA sumoylation.

**Figure 4.**
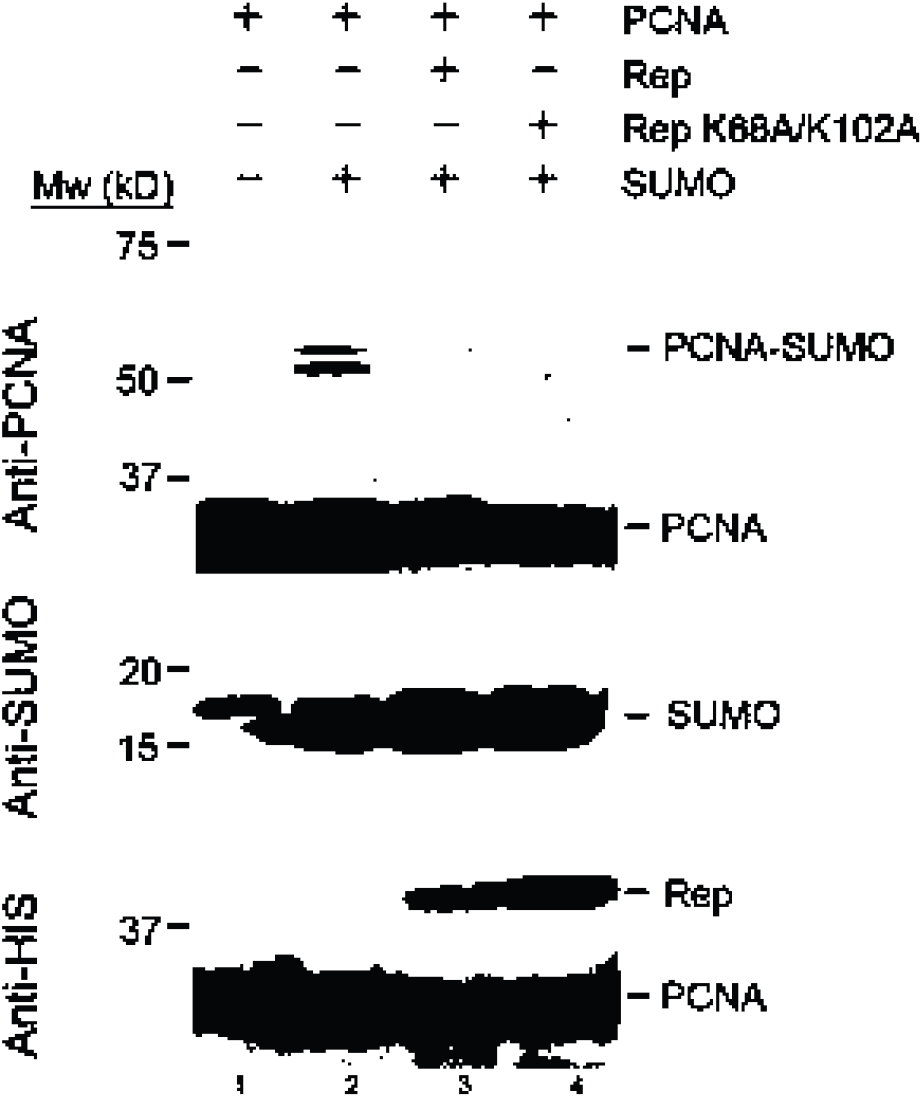
The Rep-SCE1 interaction is not essential to suppress PCNA sumoylation in bacteria. Assay is similar to Fig. 1, except that a Rep variant was co-expressed that fails to interact with SCE1 (K68A/K102A; lane 4).

The residues of PCNA that contribute to its interaction with Rep seem to spread across the protein, while the residues of Rep important for the interaction with PCNA appear to be concentrated in the middle part of this viral protein, spanning the residues 120-183 (54). To further investigate the mechanism of interference, we carried out sumoylation assays expressing truncated versions of Rep. Ten different Rep truncations were generated and expressed from the same promoter as wild-type PCNA (Fig. S2A). Expression of all these Rep mutants could be successfully detected in protein extracts from *E. coli* by western blotting using an anti-Rep antibody, except for Rep_1-68_, Rep_1-99_ and Rep_120-140_. These latter three truncations were expressed at low or undetectable levels and therefore excluded from our sumoylation assays (Fig. S2C). As described above, the expression of the full Rep protein interfered with PCNA sumoylation, producing a clear decrease in the intensity of the two sumoylated PCNA bands (Fig. S2B, lane 3). This reduction in intensity for both SUMO-modified PCNA bands was also seen when any of the Rep truncated proteins were expressed, but only in the presence of Rep_1-184_ and Rep_120-352_ the decrease in intensity of the lower band was equivalent to the effect seen with intact Rep protein (Fig. S2B, lanes 6 and 8).

### Rep interferes with PCNA sumoylation in an *Arabidopsis* sumoylation system reconstituted *in bacteria*

Analyses performed *in bacteria* with *Arabidopsis* PCNA1 by Strzalka and coworkers detected sumoylation on K254, but did not detect sumoylation of the conserved K164 (52). However, our results imply in the case of tomato PCNA that both residues can be sumoylated with a similar efficiency (Fig. 3B). The respective experimental approaches differ not only in the PCNA homologue used, but also in the origin of the sumoylation enzymes used. We used an assay previously employed to identify SUMO targets in *Arabidopsis* proteins (63) originally developed by (56), that expresses mammalian E1 and E2 enzymes and the human SUMO1. However, the results with *Arabidopsis* PCNA obtained by Strzalka and co-workers are based on a reconstituted sumoylation pathway that is entirely composed of *Arabidopsis* proteins. To determine if the discrepancy between these K164 sumoylation results is due to the disparate origin of the enzymes used, we redesigned the Mencia and Lorenzo assay replacing the mammalian E1 and E2 enzymes for their *Arabidopsis* counterparts (AtSAE and AtSCE1). Using this system, we first determined if tomato PCNA could be SUMO-modified using the *Arabidopsis* isoforms AtSUMO1, AtSUMO2 or AtSUMO3. As a positive control, we used human SUMO1. Total protein extracts were analysed by western blot using anti-PCNA or anti-His antibodies to confirm expression of the SUMO isoforms (Fig. 5A). Consistently with our previous sumoylation assay with mammalian enzymes (Fig. 3B), co-expression of PCNA and human SUMO1 produced a double band of approximately 55 kD, (Fig. 5A, lane 3). This double band was also observed in the protein extract from cells expressing either AtSUMO1 or −2, albeit at a slightly smaller molecular weight due to a different in sequence of the *Arabidopsis* SUMOs compared to the human SUMO1 (Fig. 5A, lanes 4, and 5). The same double band was also detected in the presence of AtSUMO3, however its intensity was noticeably reduced (Fig. 5A, lane 6), even with the amount of mature free AtSUMO3 being similar to that of AtSUMO1 or AtSUMO2 (Fig. 5A, lower panel, lanes 4, 5 and 6). These results indicate that tomato PCNA can be post-translationally modified by AtSUMO1 and AtSUMO2, but that it is a poor substrate for AtSUMO3 conjugation when using the *Arabidopsis* E1 and E2 enzymes.

**Figure 5.**
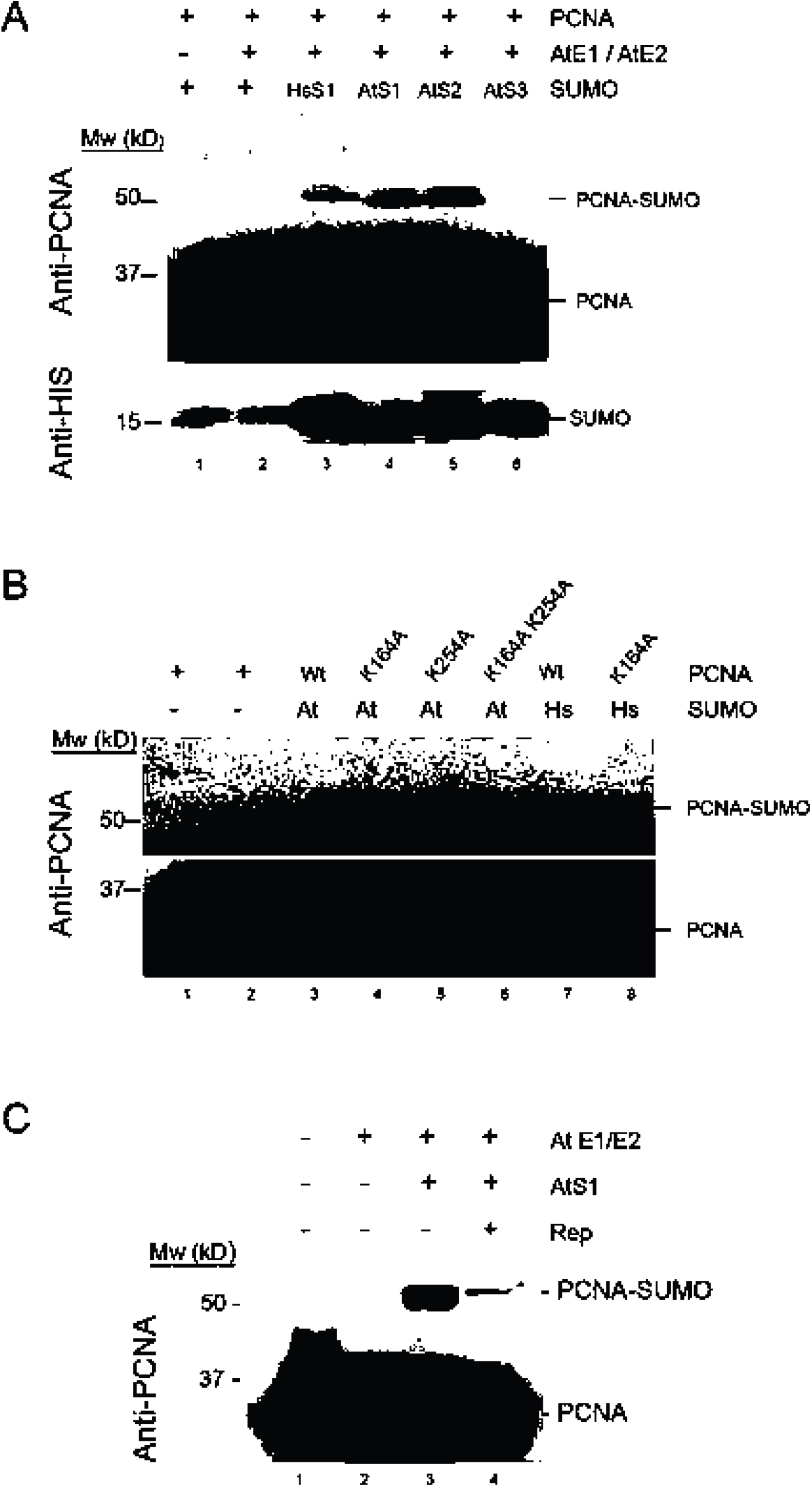
*Arabidopsis* SUMO conjugation enzymes modify the same Lys residues in tomato PCNA *in bacteria*. (A) Assay is similar to Fig. 1, except that AtE1/E2 (*Arabidopsis* SAE1, SAE2 and SCE1) were co-expressed together with the *Arabidopsis* SUMO paralogues SUMO1, −2, and −3 (AtS1/2/3/). Expression (+/-) of the individual components is indicated. E1/E2 enzymes were expressed in all samples. (B) Similar to (A), except that PCNA variants were co-expressed in which K164, K254 or both were mutated Ala. At= AtSUMO1; Hs=HsSUMO1 (lanes 4-6, 8). Wt = wildtype tomato PCNA, as used in panel A. (C) Similar to (A), except that Rep from *Tomato golden mosaic virus* (TGMV) is co-expressed (lane 4).

To confirm that the sumoylated PCNA double band corresponds to sumoylation at K164 and K254, the tomato PCNA single mutants K164A or K254A and the double mutant K164A/K254A were co-expressed in the *Arabidopsis* reconstituted assay with AtSUMO1. Expression of human SUMO1 was used as positive control. The upper band observed with wild-type PCNA disappeared when the single PCNA mutant K164A was tested (Fig. 5B, lanes 4 and 8). Expression of the single mutant K254A caused a decrease of the intensity of the lower band, while the intensity of the upper band remained unaltered when compared with wild-type PCNA (Fig. 5B, lane 5). In the case of the double mutant K164A/K254A, the upper band disappeared completely and the intensity of the lower band was greatly diminished (Fig. 5B, lane 6). These results indicate that tomato PCNA is sumoylated *in vitro* on the same lysine residues, regardless of the origin of the sumoylation machinery used.

When PCNA and Rep were expressed simultaneously from the target plasmid as a polycistronic mRNA, together with the *Arabidopsis* sumoylation machinery (AtSAE/SCE1 and AtSUMO1), the intensity of the two bands identified as mono-sumoylated PCNA was greatly reduced compared to those produced in the absence of Rep (Fig. 5C, lanes 3 and 4), confirming the result obtained using the reconstituted assay *in bacteria* (Fig. 1).

### *In planta* sumoylation of tomato PCNA is compromised by Rep

While sumoylation of plant PCNAs has been detected *in bacteria* using different reconstituted sumoylation systems, sumoylation of PCNA remains to be proven in plants cells, likely due to the low levels of modified PCNA available, probably beyond the sensitivity threshold of the detection methods used. Increasing the amount of sumoylated PCNA might allow us to infer if Rep also modulates the post-translational modification status of PCNA *in planta*. To this end, we transiently expressed Flag-tagged tomato PCNA (PCNA-Flag) together with mature AtSUMO1 in young, not fully expanded *N. benthamiana* leaves (2-week old plants), in which more cells are likely to be in the replicative stage of the cell cycle (previous studies had shown that yeast PCNA is predominantly sumoylated during the S-phase of the cell cycle (62). In addition, we subjected the harvested leaf material to a heat shock at 37ºC for 45 min to increase the levels of SUMO-conjugates (31). Western blots on PCNA protein immuno-purified with anti-Flag resin showed that PCNA accumulated in plants as monomers, dimers and trimers (of approximately 35, 70 and 100 kD, respectively) (Fig. 6A, anti-flag). These blots were also probed with anti-SUMO antibody, revealing that the PCNA-Flag monomers were also sumoylated *in planta*, giving rise to a band of approximately 50kD (Fig. 6A, anti-SUMO) equivalent to that previously detected *in bacteria* using the *Arabidopsis* system (Fig. 5). When Rep from the begomovirus *Tomato yellow leaf curl virus* was coexpressed *in planta* in the same experimental conditions, the signal for the ~50 kD band decreased, suggesting that the levels of SUMO-modified PCNA were lower when Rep was present (Fig. 6, anti-SUMO). The presence of Rep (GFP-tagged) was confirmed by immunoblotting of total protein extracts with an anti-GFP antibody (Fig. 6, anti-GFP). These results show for the first time that PCNA is sumoylated in plant cells, and confirm that Rep also compromises PCNA sumoylation *in planta*.

**Figure 6:**
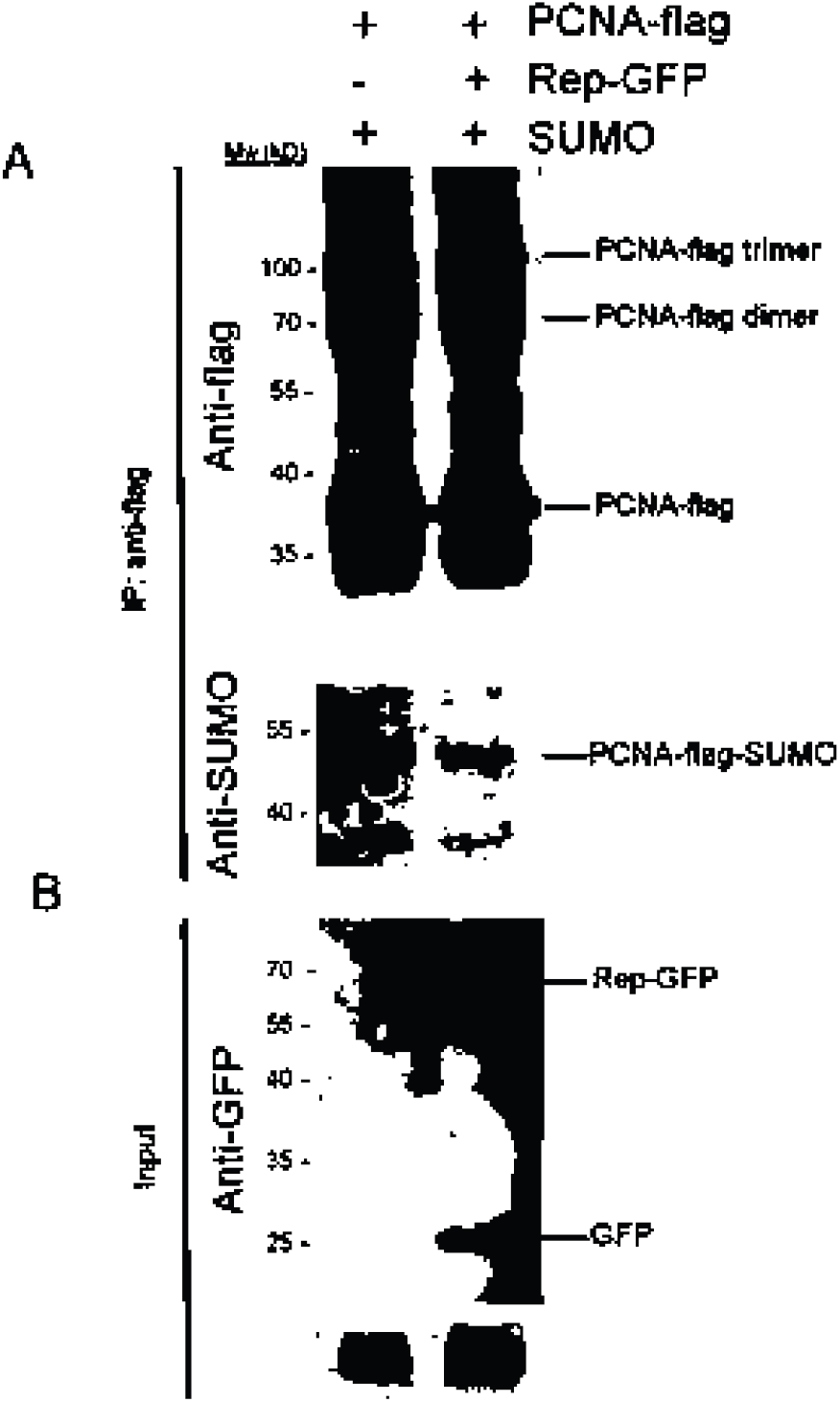
Rep from *Tomato yellow leaf curl virus* (TYLCV) compromises sumoylation of tomato PCNA *in planta*. (A) Flag-tagged tomato PCNA together with *Arabidopsis thaliana* SUMO1 was transiently expressed in *Nicotiana benthamiana* leaves in the presence or absence of GFP-tagged Rep from TYLCV. Total protein extracts were subjected to immunoprecipitation and proteins that co-eluted from the Flag affinity resin (IP:anti-Flag) were blotted with anti-Flag or anti-AtSUMO1 antibodies. The upper panel shows the enrichment of PCNA-Flag, as monomer but also as dimer and trimer (~70 and 100 kD, respectively); the anti-SUMO immunoblot reveals that in presence of Rep the ~50 kDa band, corresponding to the sumoylated monomeric form of PCNA-Flag, is reduced when Rep is coexpressed. (B) Total protein extracts (input) were analyzed by immunoblotting using anti-GFP antibody to show accumulation of Rep-GFP and by Coomassie staining to confirm equal protein loading

## DISCUSSION

PCNA is a protein highly conserved in eukaryotes that controls cell cycle regulation, DNA replication and DNA repair, through its interaction with DNA and with a plethora of proteins involved in these processes. The switching between these different PCNA functions is modulated by its post-translational modification status, mainly ubiquitination or sumoylation, which facilitates or hinders the interaction of PCNA with specific binding partners, providing a mechanism to control PCNA function (64, 65). PCNA sumoylation has a conserved role in inhibiting homologous recombination (HR) (47, 64, 65). SUMO conjugation is mediated by the sole SUMO-specific E2 enzyme SCE1. Additionally, sumoylation at K164 requires the SUMO E3 Siz1 in yeast, whereas at K127 is modified by yeast Siz2 *in vivo* but this latter modification also proceeds without an E3 *in vitro* (47, 58).

Sumoylation of the two PCNA homologues (PCNA1 and PCNA2) present in *Arabidopsis* has been described using a reconstituted sumoylation system in *E. coli* (52, 53). In the first report, sumoylation of PCNA1 was detected using *Arabidopsis* AtSUMO1 and AtSUMO3, while in the second, the authors showed efficient sumoylation of PCNA1 and PCNA2 using S. *cerevisiae* SUMO (Smt3) as well as AtSUMO1, −2, −3 and −5. In this work, we show that tomato PCNA is also sumoylated *in vitro* by human SUMO1 and AtSUMO1 and 2 (Fig. 5). However, sumoylation of PCNA with AtSUMO3 was notably inefficient compared to that obtained with the other SUMO homologues, in spite that all SUMOs were expressed in the bacteria to similar levels (Fig. 5). Taking into account that all the sumoylation enzymes used in both works were the same, this divergence must be due to the PCNA origin. The difference in sumoylation observed when using AtSUMO1/2 or AtSUMO3 is not a surprise since, like their mammalian counterparts, the *Arabidopsis* SUMO paralogues have acquired distinct expression patterns and biochemical properties (22, 39). For example, AtSUMO1/2, which are almost identical, are better substrates for conjugation than AtSUMO3 (66). The fact that S. *cerevisiae* PCNA is sumoylated when using the *Arabidopsis* enzymes (52, 67), tomato PCNA can be equally modified by mammalian as well as *Arabidopsis* SUMO-conjugating enzymes and SUMO peptides, indicates the high level of functional conservation of the mechanism to sumoylate PCNA.

Sumoylation assays of tomato PCNA identified two modification products with a similar molecular weight (Fig. 3B) that likely correspond to tomato PCNA monomers mono-sumoylated at two alternative sites (K164 versus K254). Such phenomenon was previously described for yeast PCNA, which migrated on SDS-PAGE electrophoresis with a different apparent molecular mass when sumoylated at K127 or K124 (47, 62). In none of the experiments *in bacteria* using the *Arabidopsis* or mammalian sumoylation enzymes, we have identified bands corresponding to tomato PCNA monomers simultaneously modified with two SUMO molecules (PCNA-2xSUMO), indicating that (i) the simultaneous modification of both lysine acceptor sites of one PCNA monomer or (ii) the formation of a SUMO chain (di-SUMO) on one acceptor lysine are both more inefficient than previously suggested for *Arabidopsis* PCNA (52). Considering that tomato PCNA could be efficiently sumoylated at each of two lysine acceptor residues in an alternate fashion, the lack of PCNA molecules attached to two SUMO peptides simultaneously could suggest that some hierarchy is established in the sumoylation of the lysines. Whether inhibition of the consecutive modification of both lysines with SUMO was (i) due to the biochemical characteristics of the sumoylation systems when expressed *in bacteria* or (ii) the absence of additional components such as SUMO E3 ligases, remains to be analysed. Interestingly, previous data showed that, *in bacteria*, the expression of the *Arabidopsis* SUMO E3, AtSIZ1, does not impact the sumoylation of *Arabidopsis* or S. *cerevisiae* PCNA isoforms (52, 67).

In recent years several analyses have been carried out *in planta* to identify plant SUMO targets (24, 32, 63, 68–71). The dynamic nature of the sumoylation pathway, where SUMO proteases play a major regulatory role to determine the fraction of a protein which remains sumoylated, represents a technical challenge to identify proteins modified by SUMO. In fact, PCNA sumoylation has not been detected in any of the above-mentioned studies, and even the use of plant material containing a large proportion of dividing cells as PCNA source, or transiently expressing PCNA in leaves, has failed to prove PCNA modification *in planta* (52). In this work, we show for the first time that plant PCNA, as its animal and yeast homologues, is indeed sumoylated in plant cells. The determinant use of a heat-shock to increase the accumulation of SUMO (31) allowed us to detect the sumoylation of a PCNA monomer when transiently expressing all proteins in *N. benthamiana* leaves. Interestingly, labelling of PCNA at the C-terminus with a Flag epitope does seemingly not interfere with its ability to interact with itself forming dimers or trimers.

The analysis of PCNA containing point mutations in lysine residues selected for their likelihood to be sumoylated allowed us to show that, *in bacteria*, tomato PCNA is preferentially sumoylated at two residues that are conserved in all eukaryotic PCNAs, K164, a residue reported to be sumoylated in yeast and animal, and K254. Although both residues are at the surface of the PCNA ring, they are located at opposite sides of the PCNA ring: K164 is at the back side, while K254 is at the front side of this ring. The weak band detected in the assays with the double mutant K164A/K254A with a molecular weight intermediate between the double band detected with wild-type PCNA suggests that, in the absence of those two residues, another yet unidentified lysine can be sumoylated. Whether or not that corresponds to a true third SUMO acceptor site or its modification is an artefact caused by the absence of the other two sites remains to be clarified.

Previous work with *Arabidopsis* PCNA identified K254 as one of the residues sumoylated *in bacteria*, yet failed to detect sumoylation at K164 (52). Although this study used a reconstituted system consisting of *Arabidopsis* proteins similar to the one used here, there are some experimental differences that could explain this apparent discrepancy. Mainly, to identify SUMO acceptor sites, Strzalka and co-workers used *Arabidopsis* AtSUMO3, while we used AtSUMO1. So, it could be possible that K164 is sumoylated only by AtSUMO1, while K254 can be modified by both AtSUMO1 and AtSUMO3. However, AtSUMO3 is an unique SUMO paralogue that it is only present in a small clade of the *Brassicaceae* (22). In addition, in order to analyse sumoylation Strzalka and coworkers generated *Arabidopsis* PCNA mutants replacing all but one of the lysine residues by arginines, while in our approach we only replaced those residues already proven to be sumoylated. So, we cannot rule out the possibility that in method of Strzalka and co-workers, the overall structure of PCNA is changed, which would interfere with the access of the sumoylation enzymes to specific residues. In fact, their observation that five additional lysines (located in the inner circle of the PCNA ring) can serve as SUMO acceptor site, suggests that in the absence of the main acceptor sites SCE1 will accept any available lysine as substrate. Such potentially unbiased sumoylation was previously observed for FoxM1 (72). Furthermore, overexpression of the sumoylation machinery in the presence of a target with only one lysine residue could result in the generation of false positives, given that the stoichiometric conditions on such reaction are bound to be far from physiological. However, we cannot fully exclude that *Arabidopsis* and tomato PCNA could be partially sumoylated at different residues.

Replication of the geminiviral genome fully relies on the host DNA replication machinery, including PCNA and DNA polymerases. PCNA is essential for viral replication (73) and expression of the corresponding gene is induced by the presence of Rep (74–76). Our results prove that Rep, besides binding to the PCNA protein, also interferes with its sumoylation in all modifiable lysine residues. This inhibitory effect of Rep was unique for PCNA, as it did not affect SUMO attachment to the plant protein used as a control (CAT3). The fact that PCNA sumoylation is also compromised when a Rep mutant is used that cannot interact with the SUMO conjugating enzyme SCE1, suggests that reduced SUMO conjugation of PCNA is not due to Rep inhibiting SCE1 enzymatic activity, a mechanism previously described for the Gam1 protein of the CELO virus (77). The results obtained when expressing the truncated forms of Rep point to the PCNA-binding domain in Rep as a determining factor that results in suppression of PCNA sumoylation. The results obtained by Bagewadi and colleagues (54), showing that Rep interacts with residues located all over the PCNA molecule, could indicate the reduction of PCNA sumoylation is a consequence of steric hindrance of SCE1 once Rep is bound to PCNA. This scenario would fit with the model described by Mayanagi and colleagues, where one PCNA interactor can block access of other interactors to the PCNA molecule; steric hindrance would thus fine-tune and modulate PCNA function (78–80). Specific mutants of PCNA in which the Rep-PCNA interaction is lost, will be required to confirm this hypothesis.

Sumoylation of PCNA is high in particular during the S-phase, where it would be involved in suppressing undesired recombination events between newly synthesized DNA molecules during normal fork progression. In yeast, PCNA sumoylation recruits the DNA helicase Srs2 that inhibits recombination by removing RAD51 from ssDNA, thereby disassembling an essential recombination intermediate structure [reviewed in (81)]. During replication, Srs2 aids in the repair of gaps by preventing ssDNA from being used to initiate recombination. Deletion of Srs2 or mutations of PCNA that impair its sumoylation, cause increased levels of HR (62, 82–84). The mechanisms mediated by Srs2 seem to be conserved in eukaryotes. Sumoylation of PCNA in human cells recruits PARI, a Srs2 homolog that binds to RAD51 (51, 85). In *Arabidopsis*, an Srs2 homolog was identified and shown to act as a functional DNA helicase that can process branched DNA structures that occur during synthesis-dependent strand annealing (SDSA) pathway of recombination. These properties suggest that AtSrs2 might play a role in regulating HR in plants, as predicted for its yeast homologue (86).

Several lines of evidence suggest that recombination is a key evolutionary process to generate diversity amongst ssDNA viruses (87). Recombination among geminiviral genomes has been extensively recorded for several members of the family and seems to be a consequence of a general enhancement of the recombination frequency upon infection [(88–93), reviewed in (94)] Besides the high level of recombination among the viral molecules, several experimental results also indicate that geminiviruses alter HR in plants, as infections with the begomovirus *Euphorbia mosaic virus* (EuMV) induces somatic HR events for *Arabidopsis* transgenes, specially within vein-associated tissues where this virus replicates (95).

The mechanisms of HR in ssDNA viruses remains poorly defined, but are most probably strongly influenced by the ways in which these viruses replicate. Geminiviruses replicate their circular ssDNA by three modes of action: complementary strand replication (CSR), rolling-circle replication (RCR) and recombination-dependent replication (RDR) (96, 97). It has been suggested that RDR is a replication system by which host recombination factors are utilized for geminiviral amplification and also lead to enhanced host DNA recombination (95). Rep, the only viral protein that is essential for replication, is likely to have a key role in the recruitment and assembly of this viral replisome, a protein-DNA complex that includes both viral proteins and host factors involved in DNA replication and repair, including those for HR [reviewed by (3)]. Besides its interaction with PCNA, Rep interacts with a variety of proteins involved in replication and/or HR processes such as RFC (98), RPA32 (99), Rad54 (100) or Rad51 (101). The relevance of this HR replication mechanism for geminiviruses has been highlighted recently by infecting an *Arabidopsis RAD51D* mutant with the bipartite geminivirus *Euphorbia yellow mosaic virus* (102). The results obtained showed that RAD51 D promotes viral replication at the early stages of the infection, and its presence is required for geminiviral recombination, since in the absence of RAD51D a significant decrease of both intra - and intermolecular recombinant molecules between the two DNA components of the bipartite geminivirus was observed. The fact that, in a series of experiments to develop geminivirus-based replicon for transient expression of Transcription activatorlike effector nucleases (TALENs), it was found that expression of the Rep homologue from geminivirus *Bean yellow dwarf virus* (BeYDV) increases the frequency of gene targeting (103) points to the involvement of this protein in the viral control of the HR mechanism.

Combining all our data, we propose that the interaction between Rep and PCNA modulates the protein modification status of PCNA, thus switching its cellular function to create an environment suitable for viral replication. Further, we propose that the specific reduction of PCNA sumoylation, caused by the action of Rep, is a key step to induce HR in both infecting geminiviral genomes and the genome of the infected plant cells. We propose that during viral replication Rep will interfere, possibly by its interaction with PCNA, with the ability of SCE1 to attach SUMO to PCNA. As a consequence, Srs2 binding to PCNA would be reduced, thus allowing maintenance of the Rad51-ssDNA nucleoprotein filaments generated from exposed ssDNA, which in turn will cause an increase in the level of HR recombination. Beside its effect on increasing geminiviral recombination, a higher HR activity could also have an effect on the geminiviral replication efficiency, since HR provides a mechanism for tolerating lesions that block the progression of replication forks.

## Material And Methods

### General methods

Manipulations of *Escherichia coli* strains and nucleic acids were performed according to standard methods (104). *E. coli* strain DH5α was used for subcloning. All the PCR-amplified fragments cloned in this work were fully sequenced. *E. coli* NCM631 strain was used for the sumoylation assays.

### Plasmids and cloning

Supplementary Table 1 summarises the engineering of the plasmids used in this work. Primers used in this work are summarized in Supplementary Table 2. Molecular graphics of tomato PCNA were made with the UCSF Chimera package (105) using human PCNA (protein data bank 1AXC) as reference. Chimera is developed by the Resource for Biocomputing, Visualization, and Informatics at the University of California, San Francisco (supported by NIGMS P41-GM103311).

### *In bacteria* sumoylation assays

PCNA from *Solanum lycopersicum* (obtained from pCNACT2, (55) and Rep from *Tomato golden mosaic virus* (TGMV) (positions 2461 through 2588 to nucleotide 1416; Genbank ID NC001507) PCR amplified from pDBRepTG (55) were subcloned into pET28b (Novagen, EMD Millipore, Billerica, Massachusetts) to obtain the pET28-SlPCNA and pET28-Rep plasmid, respectively. A fragment from pET28-Rep containing the RBS (ribosomal binding site) and the Rep ORF fused to a 6XHis tag was subcloned into pET28-SlPCNA, generating the pET28-SlPCNA-Rep, a vector able to produce the polycistronic mRNA for the SlPCNA and Rep proteins.

PCNA mutants were obtained using the vector pET28-SlPCNA as template and Quikchange Lightning Site-Directed Mutagenesis Kit (Stratagene, Agilent, Santa Clara, California).

To express *Arabidopsis* Catalase3 (AtCAT3) and Rep from TGMV from a polycistronic RNA, restriction fragment from pET28-Rep, containing the RBS and the Rep ORF fused to a 6XHis tag, was subcloned into pGEX-AtCAT3 to yield pGEX-AtCAT3-Rep. pGEX-AtCAT3 express the Catalase C-terminal fragment (AtCAT3Ct −419/472) fused to GST (glutathione transferase) and Flag (66).

To express C2 and SlPCNA from a polycistronic, a restriction fragment of pET28-C2 containing the RBS and the C2 ORF fused to a 6XHis tag was subcloned into pET28-SlPCNA yielding pET28-SlPCNA-C2. Previously, pET28-C2 was generated by PCR amplifying and cloning the C2 ORF of *Tomato yellow leaf curl Sardinia virus* (TYLCSV) (positions 1631 to nucleotide 1224, Genbank ID: L27708) in *Eco*RI/*Xho*I of pET-28b.

To express Rep K68A/K102A, Rep ORF was PCR amplified from pGBAL1-K68A/K102A (46) and subcloned into pET28b yielding pET28-RepK68A/K102A. A restriction fragment from pET28-RepK68A/K102A, containing the RBS and the Rep ORF fused to a 6XHis tag was subcloned downstream of the PCNA ORF in pET28-SlPCNA to yield pET28-SlPCNA-RepK68A/K102A

Truncation constructs of Rep were constructed using PCR with specific primers (see Supplementary Table 2). For the N-terminal truncations PCR fragments containing the truncated Rep were cloned in pET28b to obtain pET28-Rep_120-184_, pET28-Rep_56-130_, pET28-Rep_68-352_, pET128-Rep_120-352_, and pET28-Rep_120-352_. To express truncated Rep and SlPCNA, PCR amplified fragments of these plasmids or pET28-Rep, containing the RBS and the truncated Rep fused to a 6XHis tag were subcloned into pET28-SlPCNA to yield: pET28-SlPCNA-Rep_120-184_ pET28-SlPCNA-Rep_120-184_, pET28-SlPCNA-Rep_56-130_, pET28-SlPCNA-Rep_68-352_, pET28-SlPCNA-Rep_120-352_, and pET28-SlPCNA-Rep_120-352_, pET28-SlPCNA-Rep_1-68_, pET28-SlPCNA-Rep_1-81_, pET28-SlPCNA-Rep_1-99_, pET28-SlPCNA-Rep_1-120_ and pET28-SlPCNA-Rep_1-184_.

The polycistronic construct expressing *Arabidopsis* sumoylation E1 and E2 enzymes was generated as follows. AtSAE1, AtSAE2 and AtSCE1 were amplified from *Arabidopsis* Columbia-0 cDNA and cloned in pET28b in order to set them downstream of a RBS, yielding the plasmids pET28-AtSCE1, pET28-AtSAE1, and pET28-AtSAE2. Next, a restriction fragment of pET28-AtSCE1 was cloned into pET28-AtSAE2 to obtain pETSS and a restriction fragment of pET28-AtSAE1 was cloned into pETSS to obtain pETSS1a. Finally, to transfer the polycistronic construct to a vector with P15A ori and chloramphenicol resistance, compatible with the other plasmids used in the sumoylation assays in *E. coli*, a *Sph*I/*Eag*I fragment of pETSS1a was cloned into the *Sph*I/*Eag*I sites of pACYC184 (106) to yield pASS1a.

The CDS corresponding to the mature proteins (GG) of *Arabidopsis* AtSUMO1, 2, and 3 were PCR amplified from *Arabidopsis* cDNA and cloned in pET28b to obtain pET28-AtSUMO1, pET28-AtSUMO2, and pET28-AtSUMO3, respectively. Restriction fragments of these constructs, containing ORF of AtSUMOs fused to histidine tags were subcloned in pRHSUMO to substitute the human *Hs*SUMO1 ORF, obtaining pRHAtSUMO1, pRHAtSUMO2, and pRHAtSUMO3. Sumoylation assays with the mammalian enzymes were performed as previously described by (56). Plasmids expressing the potential sumoylation target proteins, the sumoylation enzymes and the human SUMO1, were sequentially transformed in *E. coli*. Expression was induced by adding 1mM IPTG to the culture medium in the exponential growth phase (OD_600_=0.6). Samples were taken 4 hours after induction and proteins were extracted as described in (56).

The mammalian and *Arabidopsis* sumoylation E1 and E2 enzymes are encoded in the plasmids pBADE12, and pASS1a respectively. Human SUMO1 and *Arabidopsis* SUMO 1, 2 and 3 are fused to his tags and expressed from plasmids pRHSUMO, pRHAtSUMO1, pRHAtSUMO2 and pRHAtSUMO3. In the sumoylation assays with pGEX-AtCAT3, human SUMO1 is expressed from pRKSUMO (56) instead of pRHSUMO.

### *In planta* sumoylation assay

For the *in planta* sumoylation assay, the ORF of *Solanum lycopersicum* PCNA (GenBank ID: NM_001247915, kindly provided by KeyGene N.V, Wageningen, The Netherlands) was amplified (Table S2) to generate a fragment containing the SlPCNA fused to a Flag-tag at its N-terminus. The fragment was subcloned in pJL-TRBO vector (107) to generate the pTRBO-PCNA-Flag plasmid. pK7FWG2 plasmid (108) containing Rep from TYLCV fused to EGFP (Rep Genbank ID: AF271234) was also kindly provided by KeyGene N.V. (referred to as pK7FWG2-Rep). The ORF of the mature *Arabidopsis* AtSUMO1 (residues 1-91) in pDONR221 (109) was introduced into the pGWB402 destination vector (110) using a Gateway LR Clonase II reaction (Thermo Fisher) to generate the pGWB402-SUMO1 vector.

The binary constructs pTRBO-PCNA-Flag, pK7FWG2-Rep and pGWB402-SUMO1 were introduced in *Agrobacterium tumefaciens* strain GV3101 (111) by electroporation. Single colonies were grown overnight until an OD_600_ of 0.8-1.5 in low salt LB medium (1% w/v Tryptone, 0.5% w/v yeast extract, 0.25% w/v NaCl, pH 7.0) supplemented with 20 μM acetosyringone and 10 mM MES (pH 5.6). Cells were collected by centrifugation and resuspended in infiltration medium (1× MS [Murashige and Skoog] salts (Duchefa), 10 mM MES pH 5.6, 2% w/v sucrose, 200 μM acetosyringone). The *A. tumefaciens* cultures were mixed at a ratio 1:1:1 and co-infiltrated (in the sample without Rep the pK7FWG2-Rep culture was replaced with a culture harbouring empty pGWB451 vector) in leaves of two-week old *N. benthamiana* plant at a final OD_600_ = 1. In addition, an *A. tumefaciens* strain carrying the pBIN61 with the P19 silencing suppressor from *Tomato busy shunt viru*s (TBSV) was added to every infiltration mixture at an OD_600_ = 0.5 at 2:1 ratio. Three days post-infiltration, the whole infiltrated leaves of *N. benthamiana* were harvested, placed in a petri dish on wet paper and heat shocked for 45 minutes while being floating in water bath set at 37°C in the dark. After this period, the leaf tissue was snap frozen in liquid nitrogen and stored till protein extraction.

Plant proteins were extracted as described by (112) For co-immunoprecipitations, 500 μl of the input was incubated with 30 μl resin antiFlag M2 affinity gel (Sigma-Aldrich; 50% slurry) at 4°C for 3 hours. Subsequently, the resin was collected by centrifugation (5,000g), washed 3 times with 0.5 ml washing buffer (50 mM Tris-HCl pH 7.5, 150 mM NaCl, 10% v/v glycerol, 10 mM EDTA, 0.15% v/v NP-40, 1 tablet of protease inhibitor cocktail (Roche)/50 ml buffer) and incubated at 4°C for 1 hour with 100 μl elution buffer (washing buffer + 3X Flag peptide [Sigma-Aldrich, catalog number F4799] 150 ng/μl). After incubation, the resin was transferred to Bio-Spin columns (Bio-Rad) and spun down at 1,000g for 1 minute. The eluate (immunoprecipitated proteins, referred to as IP:anti-Flag) and analysed by western blot. Details of antibodies used in this work are summarized in Supplementary Table 3.

## ACKNOWLEDGEMENTS

We thank María Lois (Center for Research in Agricultural Genomics (CRAG) CSIC-IRTA-UAB, Barcelona, Spain) for providing the plasmid pGEX-AtCAT3. This research was supported by a grant from the Spanish Ministerio de Ciencia y Tecnología (AGL2016-75819-C2-1-R). M.S.D. was awarded with a Predoctoral Fellowship from the Junta de Andalucía and an EMBO Short Term Fellowship (ASTF No: 240-05). B.S. was awarded with a Predoctoral Fellowship from the Spanish Ministerio de Educación y Cultura. We thank T. Nagakawa (Shimane University, Japan) for sharing plasmids. The Topsector T&U program Better Plants for Demands (grant 1409-036 to HvdB), including the partnering breeding companies, supported this work. F.M. is financially supported by Keygene N.V. (The Netherlands).

## References

1. Zerbini FM, Briddon RW, Idris A, Martin DP, Moriones E, Navas-Castillo J, Rivera-Bustamante R, Roumagnac P, Varsani A, Report I. 2017. ICTV Virus Taxonomy Profile: Geminiviridae. J Gen Virol 98:131–133. https://doi.org/10.1099/jgv.0.000738.

2. Jeske H. 2009. Geminiviruses. Curr Top Microbiol Immunol 331:185–226. https://doi.org/10.1007/978-3-540-70972-5_11.

3. Hanley-Bowdoin L, Bejarano ER, Robertson D, Mansoor S. 2013. Geminiviruses: masters at redirecting and reprogramming plant processes. Nat Rev Microbiol 11:777–788. https://doi.org/10.1038/nrmicro3117.

4. Settlage SB, See RG, Hanley-Bowdoin L. 2005. Geminivirus C3 protein: replication enhancement and protein interactions. J Virol 79:9885–9895. https://doi.org/10.1128/JVI.79.15.9885-9895.2005

5. Fontes EP, Gladfelter HJ, Schaffer RL, Petty IT, Hanley-Bowdoin L. 1994. Geminivirus replication origins have a modular organization. Plant Cell 6:405–416.

6. Fontes EP, Eagle PA, Sipe PS, Luckow VA, Hanley-Bowdoin L. 1994. Interaction between a geminivirus replication protein and origin DNA is essential for viral replication. Proc Natl Acad Sci USA 269:8459–8465.

7. Laufs J, Jupin I, David C, Schumacher S, Heyraud-Nitschke F, Gronenborn B. 1995. Geminivirus replication: genetic and biochemical characterization of Rep protein function, a review. Biochimie 77:765–773. https://doi.org/10.1016/0300-9084(96)88194-6.

8. Orozco BM, Hanley-Bowdoin L. 1996. A DNA structure is required for geminivirus replication origin function. J Virol 70:148–158.

9. Choudhury NR, Malik PS, Singh DK, Islam MN, Kaliappan K, Mukherjee SK. 2006. The oligomeric Rep protein of Mungbean yellow mosaic India virus (MYMIV) is a likely replicative helicase. Rev Diabet Stud 34:6362–6377. https://doi.org/10.1093/nar/gkl903.

10. Clérot D, Bernardi F. 2006. DNA helicase activity is associated with the replication initiator protein rep of tomato yellow leaf curl geminivirus. J Virol 80:11322–11330. https://doi.org/10.1128/JVI.00924-06.

11. Everett RD, Boutell C, Hale BG. 2013. Interplay between viruses and host sumoylation pathways. Nat Rev Microbiol 11:400–411. https://doi.org/10.1038/nrmicro3015.

12. Wilson VG. 2017. Viral Interplay with the Host Sumoylation System. Adv Exp Med Biol 963:359–388. https://doi.org/10.1007/978-3-319-50044-7_21.

13. Cubeñas-Potts C, Matunis MJ. 2013. SUMO: a multifaceted modifier of chromatin structure and function. Dev Cell 24:1–12. https://doi.org/10.1016/j.devcel.2012.11.020.

14. Gareau JR, Lima CD. 2010. The SUMO pathway: emerging mechanisms that shape specificity, conjugation and recognition. Nat Rev Mol Cell Biol 11:861–871. https://doi.org/10.1038/nrm3011.

15. Mazur MJ, van den Burg HA. 2012. Global SUMO Proteome Responses Guide Gene Regulation, mRNA Biogenesis, and Plant Stress Responses. Front Plant Sci 3:215. https://doi.org/10.3389/fpls.2012.00215.

16. Cappadocia L, Lima CD. 2017. Ubiquitin-like Protein Conjugation: Structures, Chemistry, and Mechanism. Chem Rev 118:889–918. https://doi.org/10.1021/acs.chemrev.6b00737.

17. Hendriks IA, Vertegaal ACO. 2016. A comprehensive compilation of SUMO proteomics. Nat Rev Mol Cell Biol 7:581–95. https://doi.org/10.1038/nrm.2016.81.

18. Hickey CM, Wilson NR, Hochstrasser M. 2012. Function and regulation of SUMO proteases. Nat Rev Mol Cell Bio 13:755–766. https://doi.org/10.1038/nrm3478.

19. Wilkinson KA, Henley JM. 2010. Mechanisms, regulation and consequences of protein SUMOylation. Biochem J 428:133–145. https://doi.org/10.1042/BJ20100158.

20. Yates G, Srivastava AK, Sadanandom A. 2016. SUMO proteases: uncovering the roles of deSUMOylation in plants. J Exp Bot 67:2541–2548. https://doi.org/10.1093/jxb/erw092

21. Novatchkova M, Tomanov K, Hofmann K, Stuible H-P, Bachmair A. 2012. Update on sumoylation: defining core components of the plant SUMO conjugation system by phylogenetic comparison. New Phytol 195:23–31. https://doi.org/10.1111/j.1469-8137.2012.04135.x.

22. Hammoudi V, Vlachakis G, Schranz ME, van den Burg HA. 2016. Whole-genome duplications followed by tandem duplications drive diversification of the protein modifier SUMO in Angiosperms. New Phytol 211:172–85. https://doi.org/10.1111/nph.13911.

23. Miura K, Hasegawa PM. 2010. Sumoylation and other ubiquitin-like post-translational modifications in plants. Trends Cell Biol 20:223–232. https://doi.org/10.1016Zj.tcb.2010.01.007.

24. Park HJ, Kim W-Y, Park HC, Lee SY, Bohnert HJ, Yun D-J. 2011. SUMO and SUMOylation in plants. Mol Cells 32:305–316. https://doi.org/10.1007/s10059-011-0122-7.

25. Huang L, Yang S, Zhang S, Liu M, Lai J, Qi Y, Shi S, Wang J, Wang Y, Xie Q, Yang C. 2009. The Arabidopsis SUMO E3 ligase AtMMS21, a homologue of NSE2/MMS21, regulates cell proliferation in the root. Plant J 60:666–678. https://doi.org/10.1111/j.1365-313X.2009.03992.x.

26. Ishida T, Fujiwara S, Miura K, Stacey N, Yoshimura M, Schneider K, Adachi S, Minamisawa K, Umeda M, Sugimoto K. 2009. SUMO E3 ligase HIGH PLOIDY2 regulates endocycle onset and meristem maintenance in Arabidopsis. Plant Cell 21:2284–2297. https://doi.org/10.1105/tpc.109.068072.

27. Ishida T, Yoshimura M, Miura K, Sugimoto K. 2012. MMS21/HPY2 and SIZ1, two Arabidopsis SUMO E3 ligases, have distinct functions in development. PLoS ONE 7:e46897. https://doi.org/10.1371/journal.pone.0046897.

28. Elrouby N. 2015. Analysis of Small Ubiquitin-Like Modifier (SUMO) Targets Reflects the Essential Nature of Protein SUMOylation and Provides Insight to Elucidate the Role of SUMO in Plant Development. Plant Physiol 169:1006–1017. https://doi.org/10.1104/pp.15.01014.

29. Castro PH, Tavares RM, Bejarano ER, Azevedo H. 2012. SUMO, a heavyweight player in plant abiotic stress responses. Cell Mol Life Sci 69:3269–3283. https://doi.org/10.1007/s00018-012-1094-2.

30. Castro PH, Verde N, Lourenço T, Magalhães AP, Tavares RM, Bejarano ER, Azevedo H. 2015. SIZ1-Dependent Post-Translational Modification by SUMO Modulates Sugar Signaling and Metabolism in Arabidopsis thaliana. Plant Cell Physiol 56:2297–2311. https://doi.org/10.1093/pcp/pcv149.

31. Kurepa J, Walker JM, Smalle J, Gosink MM, Davis SJ, Durham TL, Sung D-Y, Vierstra RD. 2003. The small ubiquitin-like modifier (SUMO) protein modification system in Arabidopsis. Accumulation of SUMO1 and −2 conjugates is increased by stress. J Biol Chem 278:6862–6872.

32. Miller MJ, Scalf M, Rytz TC, Hubler SL, Smith LM, Vierstra RD. 2013. Quantitative proteomics reveals factors regulating RNA biology as dynamic targets of stress-induced SUMOylation in Arabidopsis. Mol Cell Proteomics 12:449–463. https://doi.org/10.1074/mcp.M112.025056.

33. Conti L, Nelis S, Zhang C, Woodcock A, Swarup R, Galbiati M, Tonelli C, Napier R, Hedden P, Bennett M, Sadanandom A. 2014. Small Ubiquitin-like Modifier protein SUMO enables plants to control growth independently of the phytohormone gibberellin. Dev Cell 28:102–110. https://doi.org/10.1016/j.devcel.2013.12.004.

34. Hotson A, Chosed R, Shu H, Orth K, Mudgett MB. 2003. Xanthomonas type III effector XopD targets SUMO-conjugated proteins in planta. Mol Microbiol 50:377–389. https://doi.org/10.1046/j.1365-2958.2003.03730.x.

35. Kim J, Taylor K, Hotson A, Keegan M, Schmelz E, Mudgett M. 2008. XopD SUMO Protease Affects Host Transcription, Promotes Pathogen Growth, and Delays Symptom Development in Xanthomonas-Infected Tomato Leaves. Plant Cell 20:1915–29. https://doi.org/10.1105/tpc.108.058529.

36. Roden J, Eardley L, Hotson A, Cao Y, Mudgett MB. 2004. Characterization of the Xanthomonas AvrXv4 effector, a SUMO protease translocated into plant cells. Mol Plant Microbe Interact 17:633–643. https://doi.org/10.1094/MPMI.2004.17.6.633.

37. Castillo AG, Kong LJ, Hanley-Bowdoin L, Bejarano ER. 2004. Interaction between a geminivirus replication protein and the plant sumoylation system. J Virol 78:2758–2769. https://doi.org/10.1128/JVI.78.6.2758-2769.2004.

38. Hanania U, Furman-Matarasso N, Ron M, Avni A. 1999. Isolation of a novel SUMO protein from tomato that suppresses EIX-induced cell death. Plant J 19:533–541. https://doi.org/10.1046Zj.1365-313X.1999.00547.x.

39. van den Burg HA, Takken FLW. 2010. SUMO-, MAPK-, and resistance protein-signaling converge at transcription complexes that regulate plant innate immunity. Plant Signal Behav 5:1597–1601. https://doi.org/10.4161/psb.5.12.13913.

40. Hammoudi V, Fokkens L, Beerens B, Vlachakis G, Chatterjee S, Arroyo-Mateos M, Wackers PFK, Jonker MJ, van den Burg HA. 2018. The Arabidopsis SUMO E3 ligase SIZ1 mediates the temperature dependent trade-off between plant immunity and growth. PLoS Genet 14:e1007157. https://doi.org/10.1371/journal.pgen.1007157.

41. Lee J, Nam J, Park HC, Na G, Miura K, Jin JB, Yoo CY, Baek D, Kim DH, Jeong JC, Kim D, Lee SY, Salt DE, Mengiste T, Gong Q, Ma S, Bohnert HJ, Kwak S-S, Bressan RA, Hasegawa PM, Yun D-J. 2007. Salicylic acid-mediated innate immunity in Arabidopsis is regulated by SIZ1 SUMO E3 ligase. Plant J 49:79–90. https://doi.org/10.1111/j.1365-313X.2006.02947.x.

42. Wimmer P, Schreiner S. 2015. Viral Mimicry to Usurp Ubiquitin and SUMO Host Pathways. Viruses 7:4854–4872. https://doi.org/10.3390/v7092849.

43. Lowrey AJ, Cramblet W, Bentz GL. 2017. Viral manipulation of the cellular sumoylation machinery. Cell Commun Signal 15:27. https://doi.org/10.1186/s12964-017-0183-0.

44. Xiong R, Wang A. 2013. SCE1, the SUMO-conjugating enzyme in plants that interacts with NIb, the RNA-dependent RNA polymerase of Turnip mosaic virus, is required for viral infection. J Virol 87:4704–4715. https://doi.org/10.1128/JVI.02828-12.

45. Cheng X, Xiong R, Li Y, Li F, Zhou X, Wang A. 2017. Sumoylation of Turnip mosaic virusRNA Polymerase Promotes Viral Infection by Counteracting the Host NPR1-Mediated Immune Response. Plant Cell 29:508–525. https://doi.org/10.1105/tpc.16.00774.

46. Sánchez-Durán MA, Dallas MB, Ascencio-Ibañez JT, Reyes MI, Arroyo-Mateos M, Ruiz-Albert J, Hanley-Bowdoin L, Bejarano ER. 2011. Interaction between geminivirus replication protein and the SUMO-conjugating enzyme is required for viral infection. J Virol 85:9789–9800. https://doi.org/10.1128/JVI.02566-10.

47. Hoege C, Pfander B, Moldovan G-L, Pyrowolakis G, Jentsch S. 2002. RAD6-dependent DNA repair is linked to modification of PCNA by ubiquitin and SUMO. Nature 419:135–141. https://doi.org/10.1038/nature00991.

48. Leach CA, Michael WM. 2005. Ubiquitin/SUMO modification of PCNA promotes replication fork progression in Xenopus laevis egg extracts. J Cell Biol 171:947–954. https://doi.org/10.1083/jcb.200508100.

49. Arakawa H, Moldovan G-L, Saribasak H, Saribasak NN, Jentsch S, Buerstedde J-M. 2006. A role for PCNA ubiquitination in immunoglobulin hypermutation. PLoS Biol 4:e366. https://doi.org/10.1371/journal.pbio.0040366.

50. Gali H, Juhasz S, Morocz M, Hajdu I, Fatyol K, Szukacsov V, Burkovics P, Haracska L. 2012. Role of SUMO modification of human PCNA at stalled replication fork. Nucleic Acids Res 40:6049–6059. https://doi.org/10.1093/nar/gks256.

51. Moldovan G-L, Dejsuphong D, Petalcorin MIR, Hofmann K, Takeda S, Boulton SJ, D’Andrea AD. 2012. Inhibition of homologous recombination by the PCNA-interacting protein PARI. Mol Cell 45:75–86. https://doi.org/10.1016/j.molcel.2011.11.010.

52. Strzalka W, Labecki P, Bartnicki F, Aggarwal C, Rapala-Kozik M, Tani C, Tanaka K, Gabrys H. 2012. Arabidopsis thaliana proliferating cell nuclear antigen has several potential sumoylation sites. J Exp Bot 63:2971–2983. https://doi.org/10.1093/jxb/ers002.

53. Elrouby N, Coupland G. 2010. Proteome-wide screens for small ubiquitin-like modifier (SUMO) substrates identify Arabidopsis proteins implicated in diverse biological processes. Proc Natl Acad Sci USA 107:17415–17420. https://doi.org/10.1073/pnas.1005452107.

54. Bagewadi B, Chen S, Lal SK, Choudhury NR, Mukherjee SK. 2004. PCNA interacts with Indian mung bean yellow mosaic virus rep and downregulates Rep activity. J Virol 78:11890–11903. https://doi.org/10.1128/JVI.78.21.11890-11903.2004.

55. Castillo AG, Collinet D, Deret S, Kashoggi A, Bejarano ER. 2003. Dual interaction of plant PCNA with geminivirus replication accessory protein (Ren) and viral replication protein (Rep). Virology 312:381–394. https://doi.org/10.1128/JVI.78.6.2758-2769.2004.

56. Mencía M, de Lorenzo V. 2004. Functional transplantation of the sumoylation machinery into Escherichia coli. Protein Expres and Purif 37:409–418. https://doi.org/10.1016/j.pep.2004.07.001.

57. Castaño-Miquel L, Seguí J, Manrique S, Teixeira I, Carretero-Paulet L, Atencio F, Lois LM. 2013. Diversification of SUMO-activating enzyme in Arabidopsis: implications in SUMO conjugation. Mol Plant 6:1646–1660. https://doi.org/10.1093/mp/sst049.

58. Parker JL, Bucceri A, Davies AA, Heidrich K, Windecker H, Ulrich HD. 2008. SUMO modification of PCNA is controlled by DNA. EMBO J 27:2422–2431. https://doi.org/10.1038/emboj.2008.162.

59. Göhler T, Munoz IM, Rouse J, Blow JJ. 2008. PTIP/Swift is required for efficient PCNA ubiquitination in response to DNA damage. DNA Repair (Amst) 7:775–787. https://doi.org/10.1016/j.dnarep.2008.02.001.

60. Zhao Q, Xie Y, Zheng Y, Jiang S, Liu W, Mu W, Liu Z, Zhao Y, Xue Y, Ren J. 2014. GPS-SUMO: a tool for the prediction of sumoylation sites and SUMO-interaction motifs. Nucleic Acids Res 42:W325–30. https://doi.org/10.1093/nar/gku383.

61. Ren J, Gao X, Jin C, Zhu M, Wang X, Shaw A, Wen L, Yao X, Xue Y. 2009. Systematic study of protein sumoylation: Development of a site specific predictor of SUMOsp 2.0. Proteomics 9:3409–3412. https://doi.org/10.1002/pmic.200800646.

62. Pfander B, Moldovan G-L, Sacher M, Hoege C, Jentsch S. 2005. SUMO-modified PCNA recruits Srs2 to prevent recombination during S phase. Nature 436:428–433. https://doi.org/10.1038/nature03665.

63. López-Torrejón G, Guerra D, Catala R, Salinas J, del Pozo JC. 2013. Identification of SUMO targets by a novel proteomic approach in plants(F). J Integr Plant Biol 55:96–107. https://doi.org/10.1111/jipb.12012.

64. Mailand N, Gibbs-Seymour I, Bekker-Jensen S. 2013. Regulation of PCNA-protein interactions for genome stability. Nat Rev Mol Cell Bio 14:269–282. https://doi.org/10.1038/nrm3562.

65. Choe KN, Moldovan G-L. 2017. Forging Ahead through Darkness: PCNA, Still the Principal Conductor at the Replication Fork. Mol Cell 65:380–392. https://doi.org/10.1016/j.molcel.2016.12.020.

66. Castaño-Miquel L, Seguí J, Lois LM. 2011. Distinctive properties of Arabidopsis SUMO paralogues support the in vivo predominant role of AtSUMO1/2 isoforms. Biochem J 436:581–590. https://doi.org/10.1042/BJ20101446.

67. Colby T, Matthäi A, Boeckelmann A, Stuible H-P. 2006. SUMO-conjugating and SUMO-deconjugating enzymes from Arabidopsis. Plant Physiol 142:318–332. https://doi.org/10.1104/pp.106.085415.

68. Budhiraja R, Hermkes R, Müller S, Schmidt J, Colby T, Panigrahi K, Coupland G, Bachmair A. 2009. Substrates Related to Chromatin and to RNA-Dependent Processes Are Modified by Arabidopsis SUMO Isoforms That Differ in a Conserved Residue with Influence on Desumoylation. Plant Physiol 149:1529–1540. https://doi.org/10.1104/pp.108.135053.

69. Miller MJ, Barrett-Wilt GA, Hua Z, Vierstra RD. 2010. Proteomic analyses identify a diverse array of nuclear processes affected by small ubiquitin-like modifier conjugation in Arabidopsis. Proc Natl Acad Sci USA 107:16512–16517. https://doi.org/10.1073/pnas.1004181107.

70. Colignon B, Delaive E, Dieu M, Demazy C, Muhovski Y, Wallon C, Raes M, Mauro S. 2017. Proteomics analysis of the endogenous, constitutive, leaf SUMOylome. J Proteomics 150:268–280. https://doi.org/10.1016/jjprot.2016.09.012.

71. Rytz TC, Miller MJ, McLoughlin F, Augustine RC, Marshall RS, Juan Y-T, Charng Y-Y, Scalf M, Smith LM, Vierstra RD. 2018. SUMOylome Profiling Reveals a Diverse Array of Nuclear Targets Modified by the SUMO Ligase SIZ1 During Heat Stress. Plant Cell pii: tpc.00993.2017. https://doi.org/10.1105/tpc.17.00993.

72. Schimmel J, Eifler K, Sigurðsson JO, Cuijpers SAG, Hendriks IA, Verlaan-de Vries M, Kelstrup CD, Francavilla C, Medema RH, Olsen JV, Vertegaal ACO. 2014. Uncovering SUMOylation dynamics during cell-cycle progression reveals FoxM1 as a key mitotic SUMO target protein. Mol Cell 53:1053–1066. https://doi.org/10.1016/j.molcel.2014.02.001.

73. Morilla G, Castillo AG, Preiss W, Jeske H, Bejarano ER. 2006. A versatile transreplication-based system to identify cellular proteins involved in geminivirus replication. J Virol 80:3624–3633. https://doi.org/10.1128/JVI.80.7.3624-3633.2006.

74. Nagar S, Pedersen TJ, Carrick KM, Hanley-Bowdoin L, Robertson D. 1995. A geminivirus induces expression of a host DNA synthesis protein in terminally differentiated plant cells. Plant Cell 7:705–719.

75. Egelkrout EM, Robertson D, Hanley-Bowdoin L. 2001. Proliferating cell nuclear antigen transcription is repressed through an E2F consensus element and activated by geminivirus infection in mature leaves. Plant Cell 13:1437–1452. https://doi.org/10.1105/TPC.010004.

76. Egelkrout EM, Mariconti L, Settlage SB, Cella R, Robertson D, Hanley-Bowdoin L. 2002. Two E2F elements regulate the proliferating cell nuclear antigen promoter differently during leaf development. Plant Cell 14:3225–3236. https://doi.org/10.1105/tpc.006403.

77. Boggio R, Colombo R, Hay RT, Draetta GF, Chiocca S. 2004. A mechanism for inhibiting the SUMO pathway. Mol Cell 16:549–561. https://doi.org/10.1016/j.molcel.2004.11.007.

78. Mayanagi K, Kiyonari S, Saito M, Shirai T, Ishino Y, Morikawa K. 2009. Mechanism of replication machinery assembly as revealed by the DNA ligase-PCNA-DNA complex architecture. Proc Natl Acad Sci USA 106:4647–4652. https://doi.org/10.1073/pnas.0811196106.

79. Mayanagi K, Kiyonari S, Nishida H, Saito M, Kohda D, Ishino Y, Shirai T, Morikawa K. 2011. Architecture of the DNA polymerase B-proliferating cell nuclear antigen (PCNA)-DNA ternary complex. Proc Natl Acad Sci USA 108:1845–1849. https://doi.org/10.1073/pnas.1010933108.

80. Dovrat D, Stodola JL, Burgers PMJ, Aharoni A. 2014. Sequential switching of binding partners on PCNA during in vitro Okazaki fragment maturation. Proc Natl Acad Sci USA 111:14118–14123. https://doi.org/10.1073/pnas.1321349111.

81. Niu H, Klein HL. 2017. Multifunctional roles of Saccharomyces cerevisiae Srs2 protein in replication, recombination and repair. FEMS yeast Res 17. https://doi.org/10.1093/femsyr/fow111.

82. Aguilera A, Klein HL. 1988. Genetic control of intrachromosomal recombination in Saccharomyces cerevisiae. I. Isolation and genetic characterization of hyper-recombination mutations. Genetics 119:779–790. https://doi.org/10.1002/bies.950170210

83. Papouli E, Chen S, Davies AA, Huttner D, Krejci L, Sung P, Ulrich HD. 2005. Crosstalk between SUMO and ubiquitin on PCNA is mediated by recruitment of the helicase Srs2p. Mol Cell 19:123–133. https://doi.org/10.1016/j.molcel.2005.06.001.

84. Burgess RC, Lisby M, Altmannova V, Krejci L, Sung P, Rothstein R. 2009. Localization of recombination proteins and Srs2 reveals anti-recombinase function in vivo. J Cell Biol 185:969–981. https://doi.org/10.1083/jcb.200810055.

85. Burkovics P, Dome L, Juhasz S, Altmannova V, Sebesta M, Pacesa M, Fugger K, Sorensen CS, Lee MYWT, Haracska L, Krejci L. 2016. The PCNA-associated protein PARI negatively regulates homologous recombination via the inhibition of DNA repair synthesis. Nucleic Acids Res 44:3176–3189. https://doi.org/10.1093/nar/gkw024.

86. Blanck S, Kobbe D, Hartung F, Fengler K, Focke M, Puchta H. 2009. A SRS2 homolog from Arabidopsis thaliana disrupts recombinogenic DNA intermediates and facilitates single strand annealing. Nucleic Acids Res 37:7163–7176. https://doi.org/10.1093/nar/gkp753.

87. Martin DP, Biagini P, Lefeuvre P, Golden M, Roumagnac P, Varsani A. 2011. Recombination in eukaryotic single stranded DNA viruses. Viruses 3:1699–1738. https://doi.org/10.3390/v3091699.

88. Lefeuvre P, Martin DP, Hoareau M, Naze F, Delatte, Thierry M, Varsani A, Becker N, Reynaud B, Lett J-M. 2007. Begomovirus “melting pot” in the south-west Indian Ocean islands: molecular diversity and evolution through recombination. J Gen Virol 88:3458–3468. https://doi.org/10.1099/vir0.83252-0.

89. Padidam M, Sawyer S, Fauquet CM. 1999. Possible emergence of new geminiviruses by frequent recombination. Virology 265:218–225. https://doi.org/10.1006/viro.1999.0056.

90. Rocha CS, Castillo-Urquiza GP, Lima ATM, Silva FN, Xavier CAD, Hora Junior BT, Beserra-Júnior JEA, Malta AWO, Martin DP, Varsani A, Alfenas-Zerbini P, Mizubuti ESG, Zerbini FM. 2013. Brazilian begomovirus populations are highly recombinant, rapidly evolving, and segregated based on geographical location. J Virol 87:5784–5799. https://doi.org/10.1128/JVI.00155-13.

91. van der Walt E, Rybicki EP, Varsani A, Polston JE, Billharz R, Donaldson L, Monjane AL, Martin DP. 2009. Rapid host adaptation by extensive recombination. J Gen Virol 90:734–746. https://doi.org/10.1099/vir0.007724-0.

92. Varsani A, Shepherd DN, Monjane AL, Owor BE, Erdmann JB, Rybicki EP, Peterschmitt M, Briddon RW, Markham PG, Oluwafemi S, Windram OP, Lefeuvre P, Lett J-M, Martin DP. 2008. Recombination, decreased host specificity and increased mobility may have driven the emergence of maize streak virus as an agricultural pathogen. J Gen Virol 89:2063–2074. https://doi.org/10.1099/vir0.2008/003590-0.

93. García-Andrés S, Tomás DM, Sánchez-Campos S, Navas-Castillo J, Moriones E. 2007. Frequent occurrence of recombinants in mixed infections of tomato yellow leaf curl disease-associated begomoviruses. Virology 365:210–219. https://doi.org/10.1016/j.virol.2007.03.045.

94. Lefeuvre P, Moriones E. 2015. Recombination as a motor of host switches and virus emergence: geminiviruses as case studies. Curr Opin Virol 10:14–19. https://doi.org/10.1016/j.coviro.2014.12.005.

95. Richter KS, Kleinow T, Jeske H. 2014. Somatic homologous recombination in plants is promoted by a geminivirus in a tissue selective manner. Virology 452-453:287–296.

96. Jeske H, Lütgemeier M, Preiss W. 2001. DNA forms indicate rolling circle and recombination-dependent replication of Abutilon mosaic virus. EMBO J 20:6158–6167. https://doi.org/10.1093/emboj/20.21.6158.

97. Preiss W, Jeske H. 2003. Multitasking in replication is common among geminiviruses. J Virol 77:2972–2980. https://doi.org/10.1128/JVI.77.5.2972-2980.2003

98. Luque A, Sanz-Burgos AP, Ramirez-Parra E, Castellano MM, Gutierrez C. 2002. Interaction of geminivirus Rep protein with replication factor C and its potential role during geminivirus DNA replication. Virology 302:83–94. https://doi.org/10.1006/viro.2002.1599.

99. Singh DK, Islam MN, Choudhury NR, Karjee S, Mukherjee SK. 2006. The 32 kDa subunit of replication protein A (RPA) participates in the DNA replication of Mung bean yellow mosaic India virus (MYMIV) by interacting with the viral Rep protein. Nucleic Acids Res 35:755–770. https://doi.org/10.1093/nar/gkl1088.

100. Kaliappan K, Choudhury NR, Suyal G, Mukherjee SK. 2012. A novel role for RAD54: this host protein modulates geminiviral DNA replication. FASEB J 26:1142–1160. https://doi.org/10.1096/fj.11-188508.

101. Suyal G, Mukherjee SK, Choudhury NR. 2013. The host factor RAD51 is involved in mungbean yellow mosaic India virus (MYMIV) DNA replication. Arch Virol 158:1931–1941. https://doi.org/10.1007/s00705-013-1675-x.

102. Richter KS, Serra H, White CI, Jeske H. 2016. The recombination mediator RAD51D promotes geminiviral infection. Virology 493:113127. https://doi.org/10.1016/j.virol.2016.03.014.

103. Baltes NJ, Gil-Humanes J, Cermak T, Atkins PA, Voytas DF. 2014. DNA replicons for plant genome engineering. Plant Cell 26:151–163.

104. Burgess RC, Sebesta M, Sisakova A, Marini VP, Lisby M, Damborsky J, Klein H, Rothstein R, Krejci L. 2013. The PCNA Interaction Protein Box Sequence in Rad54 Is an Integral Part of Its ATPase Domain and Is Required for Efficient DNA Repair and Recombination. PLoS ONE. 8:e82630. https://doi.org/10.1371/journal.pone.0082630.

105. Pettersen EF, Goddard TD, Huang CC, Couch GS, Greenblatt DM, Meng EC, Ferrin TE. 2004. UCSF Chimera–a visualization system for exploratory research and analysis. J Comput Chem 25:1605–1612. https://doi.org/10.1002/jcc.20084.

106. Rose RE. 1988. The nucleotide sequence of pACYC184. Nucleic Acids Res 16:355. https://doi.org/10.1093/nar/16.1355

107. Lindbo JA. 2007. TRBO: a high-efficiency tobacco mosaic virus RNA-based overexpression vector. Plant Physiol 145:1232–1240. https://doi.org/10.1104/pp.107.106377.

108. Karimi M, Inzé D, Depicker A. 2002. GATEWAY™ vectors for Agrobacterium-mediated plant transformation. Trends Plant Sci 7:193195. https://doi.org/10.1016/S1360-1385(02)02251-3

109. Mazur MJ, Spears BJ, Djajasaputra A, van der Gragt M, Vlachakis G, Beerens B, Gassmann W, van den Burg HA. 2017. Arabidopsis TCP Transcription Factors Interact with the SUMO Conjugating Machinery in Nuclear Foci. Front Plant Sci 8:2043. https://doi.org/10.3389/fpls.2017.02043.

110. Nakamura S, Mano S, Tanaka Y, Ohnishi M, Nakamori C, Araki M, Niwa T, Nishimura M, Kaminaka H, Nakagawa T, Sato Y, Ishiguro S. 2010. Gateway binary vectors with the bialaphos resistance gene, bar, as a selection marker for plant transformation. Biosci Biotechnol Biochem 74:1315–1319. https://doi.org/10.1271/bbb.100184.

111. Koncz C, Schell J. 1986. The promoter of TDNA gene controls the tissue-specific expression of chimaeric genes carried by a novel type of binary vector. Molec Gen Genet 204:383–396. https://doi.org/10.1007/BF00331014.

112. de la Paz Sánchez M, Torres A, Boniotti MB, Gutierrez C, Vázquez-Ramo JM. 2002. PCNA protein associates to Cdk-A type protein kinases in germinating maize. Plant Mol Biol 50:167–175. https://doi.org/10.1023/A:1016029001537.

